# Bio-Physical Modeling of *Galvanic* Human Body Communication in Electro-Quasistatic Regime

**DOI:** 10.1101/2020.11.23.394395

**Authors:** Nirmoy Modak, Mayukh Nath, Baibhab Chatterjee, Shovan Maity, Shreyas Sen

## Abstract

Human Body Communication (HBC) is an alternative to radio wave-based Wireless Body Area Network (WBAN) because of its low-loss, wide bandwidth leading to enhanced energy efficiency. HBC also shows better performance in terms of physical security as most of the signal is confined within the body. To obtain optimum performance and usability, modeling of the body channel plays a vital role. Out of two HBC modalities, Galvanic HBC has the promise to provide lower loss compare to Capacitive HBC for shorter channel length. In this paper, we present the first lumped element based detailed model of Galvanic HBC channel which is used to explain the dependency of channel loss on the material property of skin, fat and muscle tissue layer along with electrode size, electrode separation, geometrical position of the electrodes and return path capacitance. The model considers the impedance of skin and muscle tissue layers and the effect of various coupling capacitances between the body and Tx/Rx electrodes to the Earth-ground. A 2D planner structure is simulated in HFSS to prove the validity of the proposed model. The effect of *symmetry* and *asymmetry* at the transmitter and receiver end are also explained using the model. The experimental results show that, due to the mismatch at the transmitter and receiver side, the loss increases gradually with channel length and saturates to a finite value as channel length becomes significantly longer compare to the transmitting or receiving electrode pair separation.

## I. Introduction

HUMAN Body Communication (HBC) was first introduced by Zimmerman [1] in 1995. It is gaining significant recognition in the field of continuous health care monitoring as well as in the field of Wearable and Implantable devices which uses Body Area Network (BAN) for communication [2]–[5]. Wearable devices like ECG band, wristwatch, flexible electronics, etc. use BAN to collect the vital information continuously and generate appropriate signals whenever an anomaly is detected. Apart from this, those small and tiny sensors do not restrict body movements which allow the acquisition of biomedical signals uninterruptedly increasing the duration of operation without degrading the comfort level of a patient. High-Security level and data privacy are few additional advantages which make it more popular over regular Bluetooth network [6], [7]. In any communication method, an in-depth understanding of the channel-model is crucial for efficient circuits and systems development. While recently there has been a Bio-Physical model proposed for capacitive HBC [8], there exists no such model for Galvanic HBC considering the simultaneous effect of parasitic capacitances and body impedance. Therefore, modeling the Galvanic HBC channel is a strong open need in the field of Human Body Communication.

The coupling of the signal to the human body can be done using two different HBC modalities: (a) Capacitive HBC and (b) Galvanic HBC. In case of Capacitive HBC [Fig. 1(a)], the signal electrode of the transmitter is connected to the body with ground electrode floating [1]. The injected signal is sensed using a receiver connected somewhere else to the body in a similar fashion. The floating electrodes of the Rx-Tx pair coupled with the Earth-ground and form a return path for signal whereas the body provides the forward signal path. The path loss of Capacitive HBC is mostly determined by the return path capacitances and load capacitance due to the low impedance forward path through the body as described in [8].

**Fig. 1:**
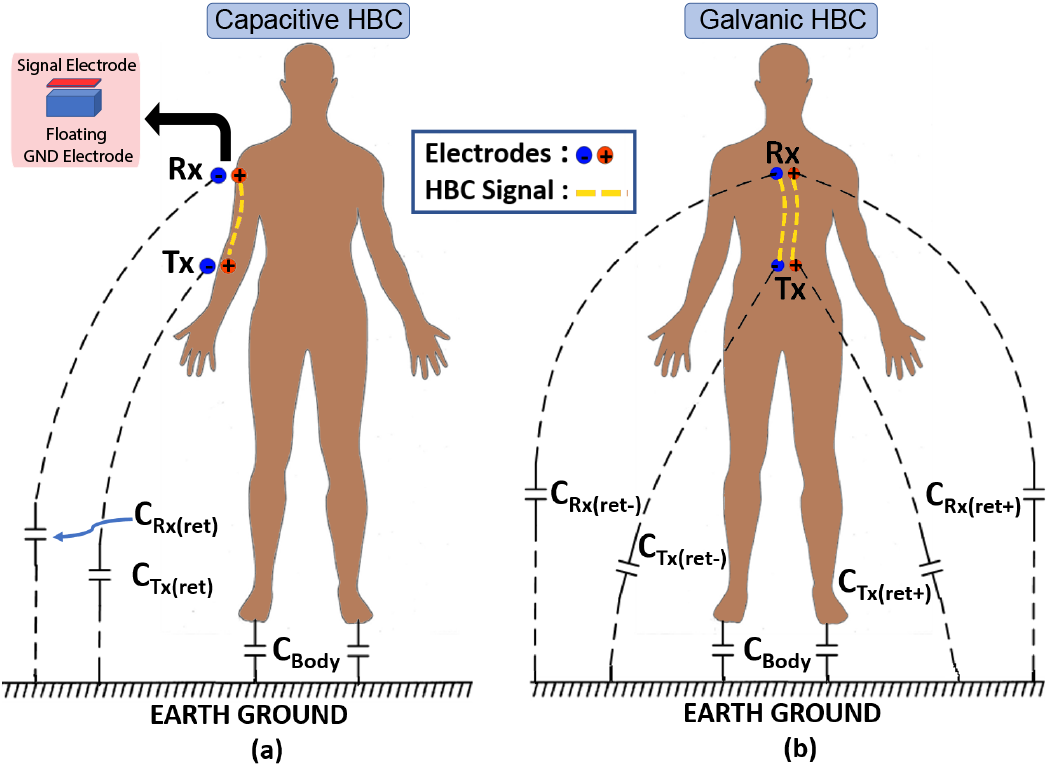
HBC coupling modalities: (a) Capacitive (b) Galvanic

Data transfer through human body using Galvanic HBC first proposed by Wegmueller [9] in 2010. Unlike Capacitive HBC, Galvanic HBC [Fig. 1(b)] uses a pair of electrodes to inject signal into the body. At the receiver side, another pair of electrodes receive the signal. Electrode size, electrode to the Earth-ground parasitic capacitance and the body channel length play a crucial role to define the overall Galvanic channel characteristics. As both the Tx-Rx electrode pairs stay in contact with body, Galvanic HBC forms a more complex return path for signal compare to the Capacitive HBC. The Galvanic return path is partially through the body and partially through the Earth-ground parasitic capacitances associated to the electrodes. For a short channel length, when the distance between the Tx and Rx is comparable to the Tx/Rx electrode pair separation, Galvanic HBC shows a distance-dependent loss, as most of the signal takes in body return path. As the signal travelling through the body medium for short distance can be considered like a dipole-dipole interaction between transmitter and receiver which is highly dependent on the dipole separation and orientation. Here, the transmitter electrode pair acts as a dipole transmitter and the receiver electrode pair acts as a dipole receiver if they are placed very close to each other. It has also been observed that in case of a longer channel length, when the distance between the Tx and Rx is significantly more compare to the Tx/Rx electrode pair separation, the Galvanic loss is determined by the mismatch present in the parasitic electrode capacitances. At longer channel length, signal taking the outer body return path encounter significantly low loss compare to the signal taking in-body return path. This is due to the weak in-body dipole-dipole interaction between the Tx and Rx which reduces with distance whereas the interaction between the Tx and Rx through outer body parasitic capacitive network is independent of channel length. Therefore, at a longer channel length, galvanic loss becomes independent of channel length, due to the Tx-Rx interaction mostly through outer body capacitive network and the loss number is determined by the mismatch present in the parasitic capacitances. It is also seen that at longer channel length due the Tx-Rx interaction through outer body capacitive network, the behaviour of the Galvanic channel becomes similar to capacitive HBC even if the excitation and termination are Galvanic.

In this paper, we proposed the first lumped element based bio-physical circuit model of Galvanic HBC considering the parameters such as thickness, conductivity, permittivity of skin, fat and muscle layers along with the return path capacitances associated to each of the electrodes. The model itself takes care of the geometric position of the electrodes which helps to understand the short channel and long channel galvanic loss in a better way. The proposed model helps to improve the accuracy in predicting the transmitted and received signal under multiple scenarios such as balanced or unbalanced input and output. During the analysis, we have considered a planner structure of thin skin and fat layer on top of a thicker muscle layer. The material properties such as dielectric constant, conductivity, loss tangent, etc. of the above-mentioned layers are considered from [10] and [11] depending on the frequency of operation. Finite Element Analysis (FEA) results and the actual HBC channel response at multiple frequencies with different channel lengths are used to validate the proposed model and its accuracy.

The rest of the paper is organized as follows: In Section II, compares some previous studies where authors try to characterize or find the channel loss in terms of body impedance and TRx position. The biophysical model of the galvanic HBC is modeled in Section III. Section IV discuss the development of Galvanic HBC model by replacing skin-fat and muscle layers by its equivalent electrical lumped-elements. Section V formulates the channel loss expression considering a balanced or symmetric scenario at the transmitter and receiver side. Section VI discusses the channel loss considering unbalanced or asymmetry at the transmitter and receiver side. Section VII extends the concept of Section V and VI to a cylindrical model which is a more realistic scenario. Section VIII discusses the measurement setup and validates the measured loss with our proposed model. Section IX shows the implementation of capacitive HBC Bio-physical model using the proposed Galvanic HBC Bio-physical model. Section X concludes the paper.

## II. Background

The concept of Galvanic HBC first introduced by Wegmueller [9] in 2010. Wegmueller shows, the coupling of the signal to the human body can be done differentially using a pair of electrodes at transmitter side and another electrode pair can be used to acquire the signal differentially at receiver side. Very few studies [12]–[15] have been carried out since then to characterize the Galvanic HBC channel and its loss. However, none of the studies explicitly explains the detailed model of the galvanic HBC channel. In [12], Wegmueller *et* al. studied the influence of the electrode size and human body joints on the channel for a stimulus input based on the position of the transmitter and receiver at different locations on the body. K. Ito and Y. Hotta in [13] proposed a four terminals circuit model for galvanic HBC and validated the model by simulation. X. M. Chen *et* al. derived the channel capacity for a general HBC channel using a water-filling algorithm and Shannon’s theorem [14]. D. Ahmed *et* al. in [15] proposed a simulation-based model for arm and found the effect of multiple skin tissue layers for galvanic coupling intra-body communication. They also discussed the effect of the bending angle on the received signal.

The studies conducted so far characterize the human body channel based on overall body impedance. The goal of this paper is to find a lumped element based circuit model that can be used to estimate the galvanic HBC channel loss. The paper also establishes the effect of the symmetry or asymmetry present at the transmitter and receiver side on galvanic loss due to the mismatch in return path capacitances using the same proposed model.

## III. Bio-Physical Behaviour of Galvanic HBC

The transmission of electrical signal around the human body happens through multiple tissue layers. The signal generated by the transmitter first gets coupled to the body through the skin and fat tissue layer and coupled signal mostly travels around the body through the muscle layer because of the high water content and interstitial fluid present in the muscle which provide a low resistive forward path. Due to the lossy dielectric nature of the skin and fat tissue layer, the coupling of the signal to the body at a higher frequency is mostly capacitive and the coupling efficiency increases as frequency increases. Whereas at low frequency, the skin and fat layer act as resistance, and the coupling of the signal to the body does not change with frequency. At that low frequency range, the signal encounters resistive division between the skin-fat and muscle layer. To prove these facts, we have considered a planner structure (200*cm* × 200*cm*) of skin-fat (1*mm* thick) and muscle (1*cm* thick) layer and simulated the model in Ansys HFSS, a Finite Element Analysis (FEA) tool. A Galvanic transmitter is placed at the center of the structure and the received voltage is measured at different distances. Three different simulation results are shown in Fig. 2. The first plot Fig. 2(a) shows the galvanic loss vs frequency at a fixed channel length (50*cm*) with electrode size as a parameter. As we can see at low frequency range the loss is constant with frequency and after a certain frequency, the loss reduces. This nature indicates that loss at lower frequency is due to the resistive division of the signal between the skin-fat and muscle layer. It is also seen that a bigger electrode helps better coupling. In the second plot Fig. 2(b), the loss vs channel length for a fixed electrode size (*r* = 1*cm*) with frequency as a parameter is shown. It is seen that higher frequency provides better coupling and overall improvement of the loss due to lower skin-fat impedance in the signal path. The sharp decay of the signal at the edge of the electrodes indicates the voltage drop between the skin-fat layer, higher the frequency lesser the drop is. In the last figure Fig. 2(c), the loss vs channel length is shown at fixed frequency (100*kHz*) for different electrode size. From the plot, it is observed that a bigger electrode provides lower loss because of the better coupling between the electrode and muscle layer, due to the presence of higher skin-fat capacitance or lower impedance along the signal path. It is also observed that, if we neglect the coupling loss, the distance dependent loss increases at a rate of −40*dB* per decade increase in channel length, which signifies the confinement of the signal within the 2D tissue layer.

**Fig. 2:**
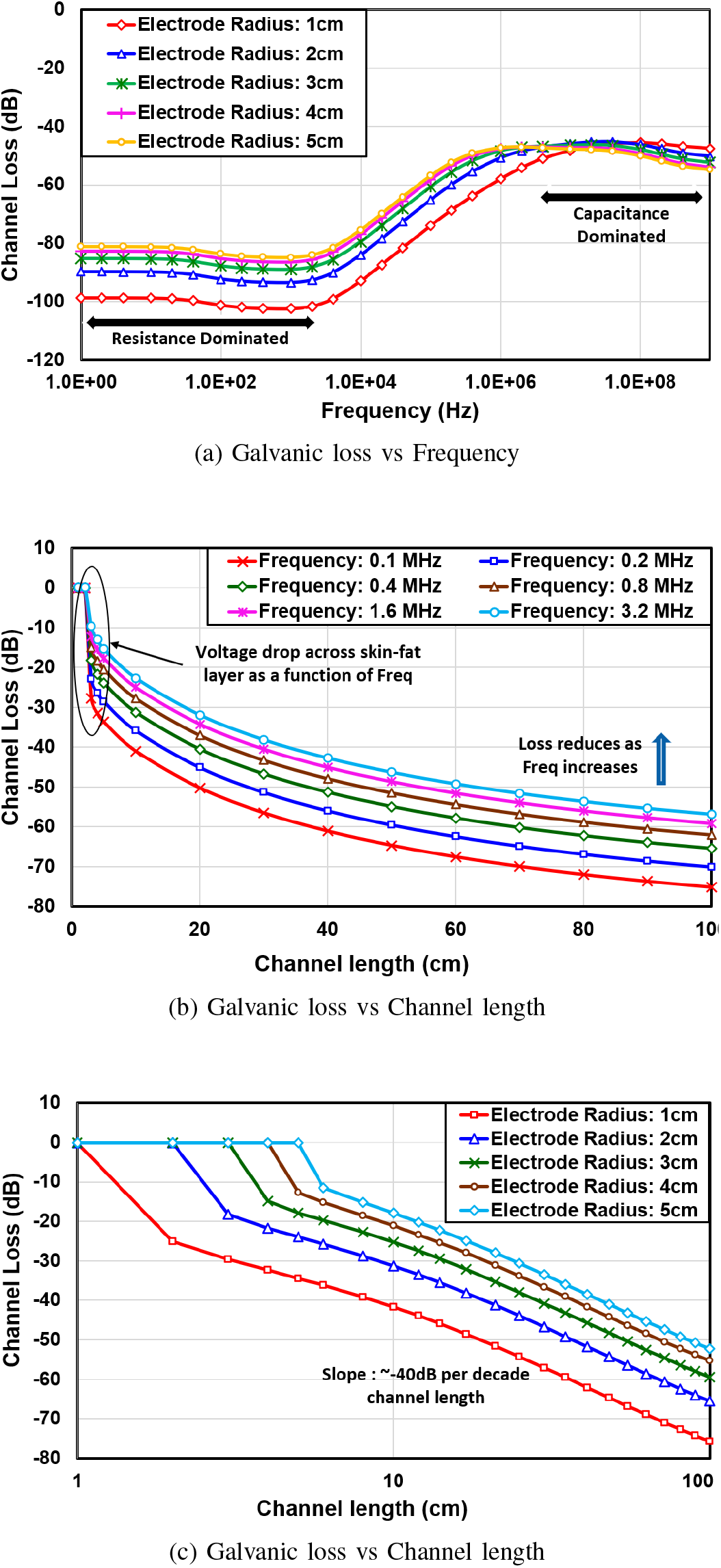
HFSS Simulation results of a 200*cm* × 200*cm* plan-ner skin-fat-muscle tissue layers under multiple scenario (a) Simulated result of Galvanic loss vs frequency for a 50cm fixed c hannel l ength w ith e lectrode s ize a s a p arameter, (b) Simulated result of Galvanic loss vs channel length for a fixed electrode size at different frequencies, (c) Simulated results of Galvanic loss vs channel length at fixed frequency for different electrode sizes.

Therefore, the HFSS simulation results show significant dependency of galvanic loss on coupling capacitances and channel length. Keeping this in mind, we are trying to model the lumped circuit capturing all the effects present in the galvanic HBC. We will also include the return path capacitance associated with each electrode to provide a better understanding of some non-ideal conditions like input/output symmetry or asymmetry in the channel.

## IV. Bio-Physical Model Development

In this section, we have shown the analysis to find the equivalent lumped circuits for the skin-fat layer, muscle layer and return path capacitance. The same 2D planner structure with skin-fat and muscle layer used earlier in HFSS simulation is used here for the analysis. Although the human body is not 2D this analysis helps to understand the behavior of the HBC channel in a simplistic way. Later on, we have extended the concept of planner structure to a more complex human body in Section VII.

### A. Equivalent Circuit for skin and fat layer

Coupling of the HBC signal to the human body is mainly affected by the thickness and the dielectric properties of the skin and fat tissue layer. In the Electro-Quasistatic region, due to the dielectric nature of the skin layer, the effective capacitance and resistance can be expressed by the Eq. 1 and Eq. 2. The resistance parallel to the capacitance is due to the lossy nature of the skin dielectric.

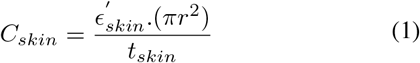

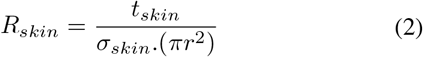

where 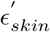 is the absolute permittivity and *σ*_*skin*_ is the conductivity of the skin tissue layer at that frequency of operation. Whereas, *t*_*skin*_ is the thickness of the skin tissue layer and *r* is the radius of the electrode connected.

Similar to the skin layer, the fat layer also can be represented using a combination of capacitor Eq. 3 and resistor Eq. 4 depending on the thickness and the dielectric properties of the fat at that frequency.

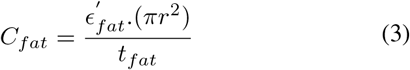

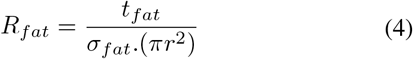

The lumped circuit representation of the skin and fat layer for a particular electrode is shown in Fig. 3. Although we have shown the equivalent circuit model of skin and fat tissue layer, the experimental data [10] [11] shows that resistance and capacitance due to the fat tissue layer are much smaller than the resistance and capacitance introduce by the skin tissue layer. Therefore if we neglect the fat tissue layer the equivalent circuit reduces to a single RC network representing the skin properties and bridged between electrode and muscle tissue layer.

**Fig. 3:**
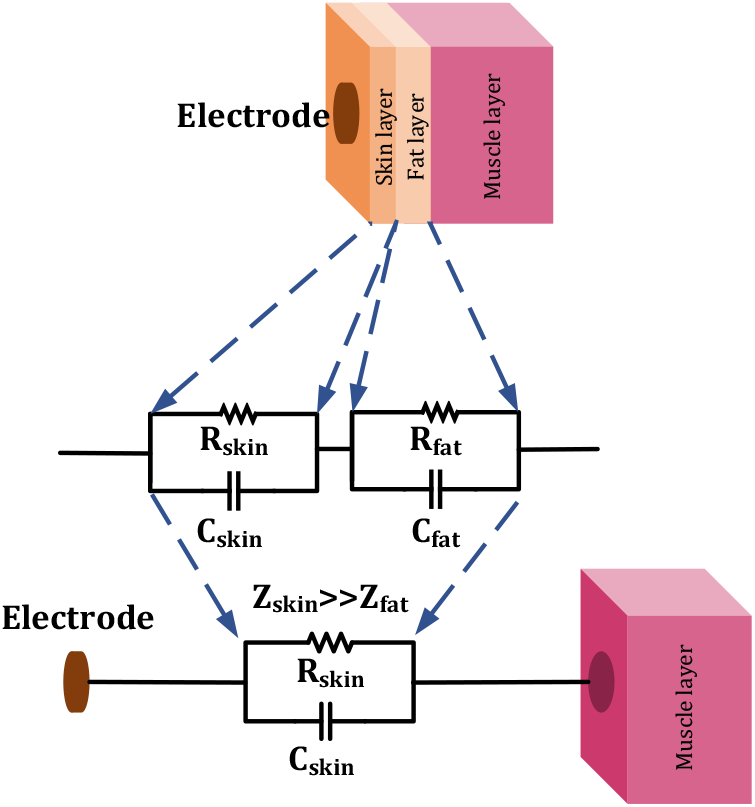
Lumped circuit equivalent of the skin-flat layer.

### B. Equivalent Circuit for Muscle layer

The transmitted current capacitively coupled to the muscle layer through the capacitance formed by the skin tissue layer which is sandwiched between the electrode and the highly conductive muscle layer. To find the current distribution in the muscle layer a planner muscle structure of finite thickness is considered as shown in Fig. 4. The portion of the muscle layer which is right below the electrode can be considered to have an *equipotential surface* due to the capacitive coupling with the electrode. For our analysis, we have considered two regions, *Region A* and *Region B* on the muscle layer which are right below the positive and negative transmitting electrodes respectively. Current *I*, which is coupled to the body through skin tissue layer is flowing from *Region A* to *Region B*. *Region C* and *Region D* are the two regions on the muscle layer, right below the negative and positive electrode of the receiver. If the lateral dimension of the muscle layer is much more than the electrode separation, the muscle tissue layer can be considered as an infinitely stretched conductor of finite thickness *h*_*muscle*_. Now, as shown in **Appendix A**, considering *Region A* and *Region B* as an input port and *Region D* and *Region C* as an output port, the entire muscle tissue layer can be represented as a two-port network where the impedance(Z) parameters of the network are as follows:

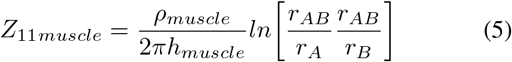

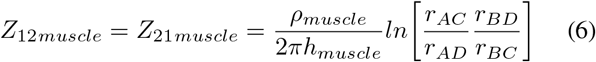

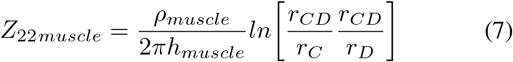

here *ρ*_*muscle*_ is the resistivity of the muscle tissue layer; *r*_*i*_ (where i=A or B or C or D) is the radius of *Region i* and *r*_*ij*_ (where i or j =A or B or C or D) is the center to center distance between *Region i* and *Region j*.

**Fig. 4:**
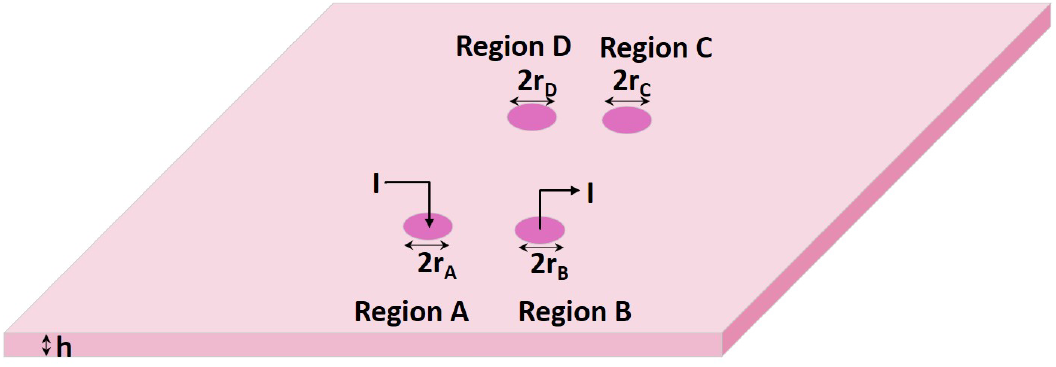
A planer structure of muscle layer with coupled current I flowing from *Region A* to *Region B* which are the regions right below the transmitting electrodes. Whereas, *Region D* and *Region C* are the regions on the muscle layer right below the receiving electrodes.

The muscle layer can also be represented using an impedance network as shown in Fig. 5 where the impedance *Z*_*T muscle*_, *Z*_*Rmuscle*_ and *Z*_0*muscle*_ are the function of the impedance parameters *Z*_11*muscle*_, *Z*_12*muscle*_ and *Z*_22*muscle*_ as described by Eq. 8, Eq. 9 and Eq. 10.

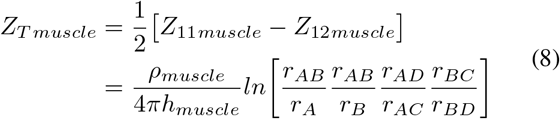

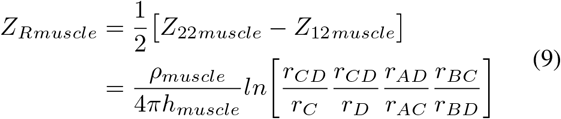

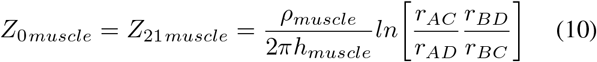

**Fig. 5:**
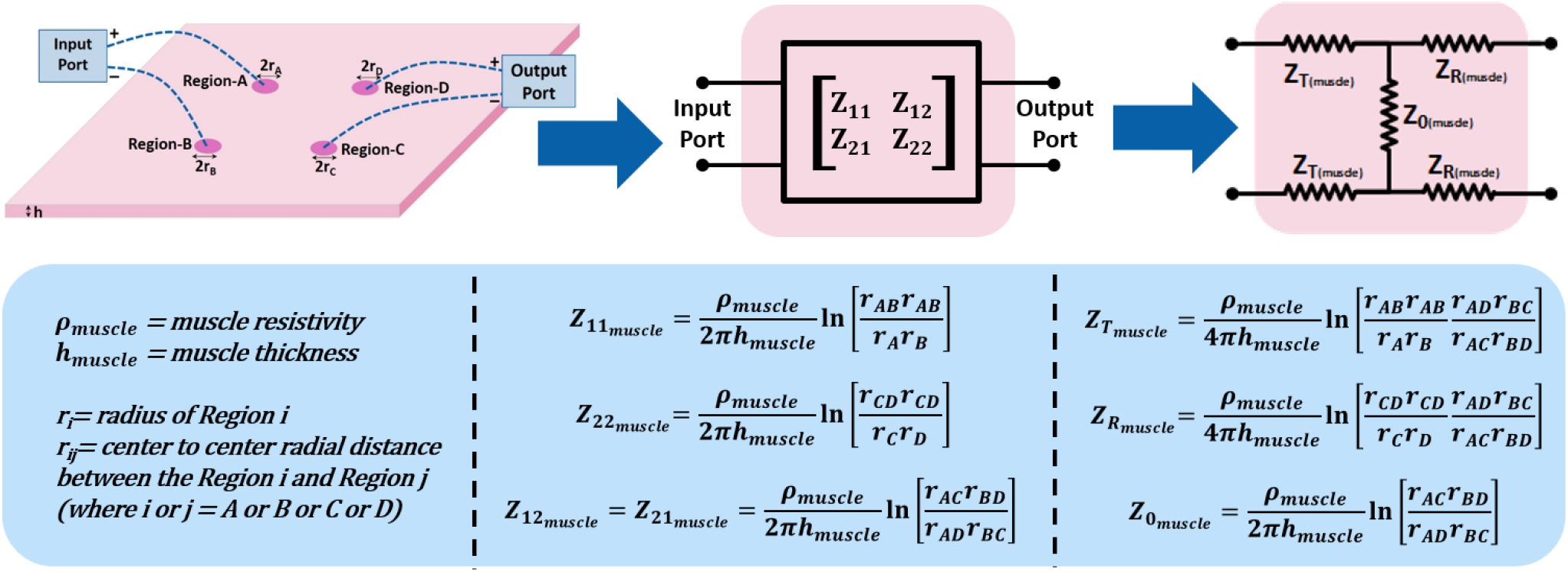
The equivalent two port representation of muscle layer with all the Z-parameters of the network representing the geometrical and electrical properties of the muscle layer as impedance parameters.

### C. Return path capacitance

In Human Body Communication, the return path capacitance plays a vital role to establish the overall channel loss. For example, in Capacitive HBC, body provides the froward path for the signal transmission and the return path for the signal is formed by the electrode to Earth-ground capacitance (return path capacitance) associated to the floating electrode of the transmitter and receiver. Return path capacitances have much higher impedance compare to the forward path body impedance. Due to the presence of high impedance return path, the overall channel loss is primarily determined by the value of return path capacitances associated to the floating electrodes [8]. Unlike Capacitive HBC, Galvanic HBC [Fig. 1(b)] uses a pair of electrodes to inject and receive signals. As both the Tx-Rx electrodes stay in contact with the body, Galvanic HBC forms a more complex return path capacitance network which affects the overall channel loss drastically. Therefore adding the proper return path capacitances to the bio-physical model is an absolute necessity and has been missing from literature.

According to the study in [16], for a circular electrode of radius *r* and thickness *h* (*h* ≪ *r*), the effective *self capacitance* (Capacitance with respect to the Earth-ground) is approximately equal to 8*ϵ*_0_*r*. Due to the body contact to the electrode, there is another electrode to Earth-Ground capacitance through body; we call it *C*_*body*_. Now, the self capacitance of the electrode, *C*_*body*_ along with any extra capacitance added to the electrode node because of transmitter or receiver connection, all together act as a return path capacitance of that electrode node. Therefore for an electrode, the value of the return path capacitance associated to it is given by the Eq. 11.

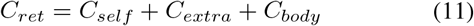

where *r* is the radius of the electrode and *C*_*extra*_ is the extra capacitance present at that electrode node due to the transmitter or receiver connection.

### D. Complete Bio-physical Model

The complete proposed biophysical model of the galvanic HBC is shown in Fig.6. Each portion of the skin tissue layer which is between the muscle layer and electrodes has been replaced using an equivalent RC network. The value of the capacitance and resistance for the skin layer is determined by the corresponding electrode area, dielectric property and thickness of the skin layer as expressed by Eq. 1 and Eq. 2. In case of muscle layer, electrical properties, thickness and the geometric position of the transmitter and the receiver electrodes determine the values of impedance *Z*_*T muscle*_, *Z*_*Rmuscle*_ and *Z*_0*muscle*_ as expressed by Eq. 8, Eq. 9 and Eq. 10 respectively. *C*_*ret*(*Tx*+)_ and *C*_*ret*(*Tx−*)_ are the return path capacitances associated to the transmitting electrodes while *C*_*ret*(*Rx*+)_ and *C*_*ret*(*Rx*−)_ are the return path capacitances associated to the receiving electrodes. The other ends of all the return path capacitances are connected to the Earth-ground which forms a signal path between the transmitter and the receiver which is outside of the human body.

**Fig. 6:**
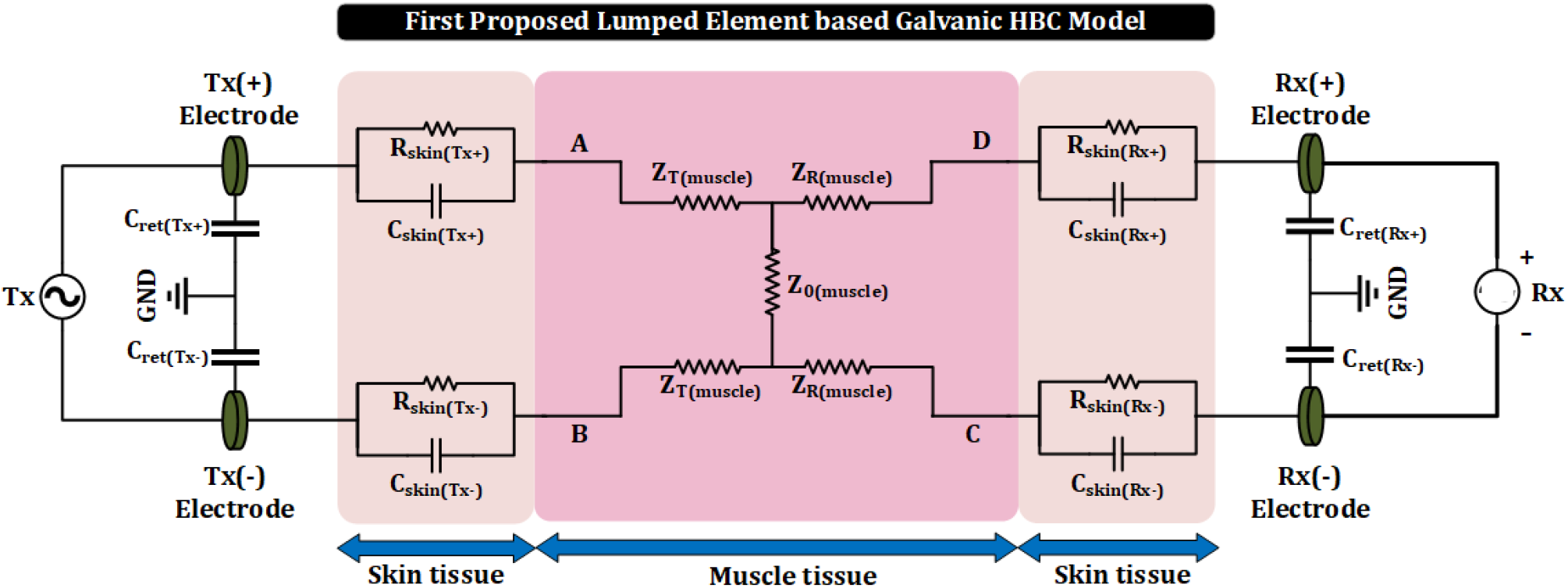
The complete bio-physical model of galvanic HBC which represents the skin and muscle tissue layer with its equivalent lumped impedance.

## V. Galvanic channel loss for symmetric or balanced channel

A symmetric or balanced condition occurs when both the pairs of transmitting and receiving electrodes possess identical size (or radius) and shape with equal return path capacitances such that *C*_*ret*(*Tx*+)_ = *C*_*ret*(*Tx*−)_ and *C*_*ret*(*Rx*+)_ = *C*_*ret*(*Rx*−)_. Balanced condition in Galvanic HBC channel provides a lot more flexibility to formulate the channel loss using a simple mathematical expression. Fig. 7 shows a balanced galvanic arrangement with channel length *d* and electrode separation *s*. Angle *θ* represents the angular position of the receiver with respect to the transmitter. Due to the identical electrodes, we can consider all the skin capacitances are equal to *C*_*skin*_ and resistances are equal to *R*_*skin*_. While the return path capacitances associated to each of the electrodes are equal to *C*_*ret*_. Using Eq. 8, Eq. 9 and Eq. 10 the impedance parameters *Z*_*T muscle*_, *Z*_*Rmuscle*_ and *Z*_0*muscle*_ of the muscle layer can be represented as a function of *s*, *d* and *θ*; such as

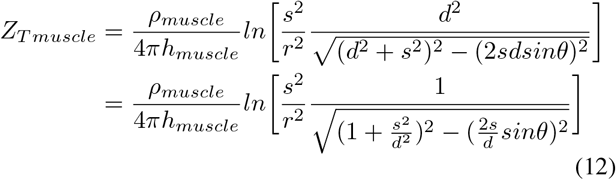

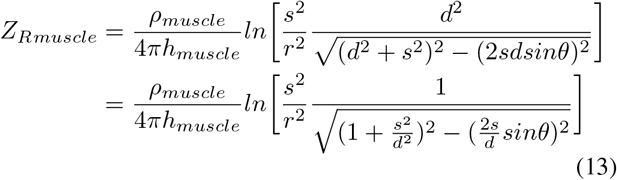

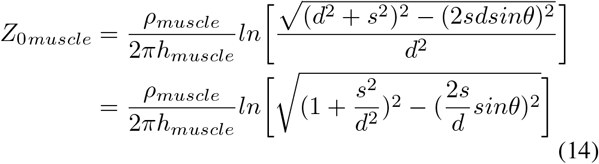

where, we have considered, electrode radius *r*_*A*_ = *r*_*B*_ = *r*_*C*_ = *r*_*D*_ = *r* ; electrode pair separation *r*_*AB*_ = *r*_*CD*_ = *s*; channel length *r*_*AD*_ = *r*_*BC*_ = *d*; 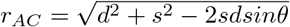 and 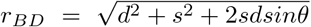. The expression of *r*_*AC*_ and *r*_*BD*_ are calculated by using the properties of triangle in trigonometry. From Eq. 12, Eq. 13 and Eq. 14 we can conclude that for a fixed *θ*, as *d* increases *Z*_*T muscle*_ and *Z*_*Rmuscle*_ increases while *Z*_0*muscle*_ decreases. Similarly, for a fixed channel length *d*, as 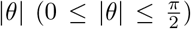 increases *Z*_*T muscle*_ and *Z*_*Rmuscle*_ increases while Z_0*muscle*_ decreases.

**Fig. 7:**
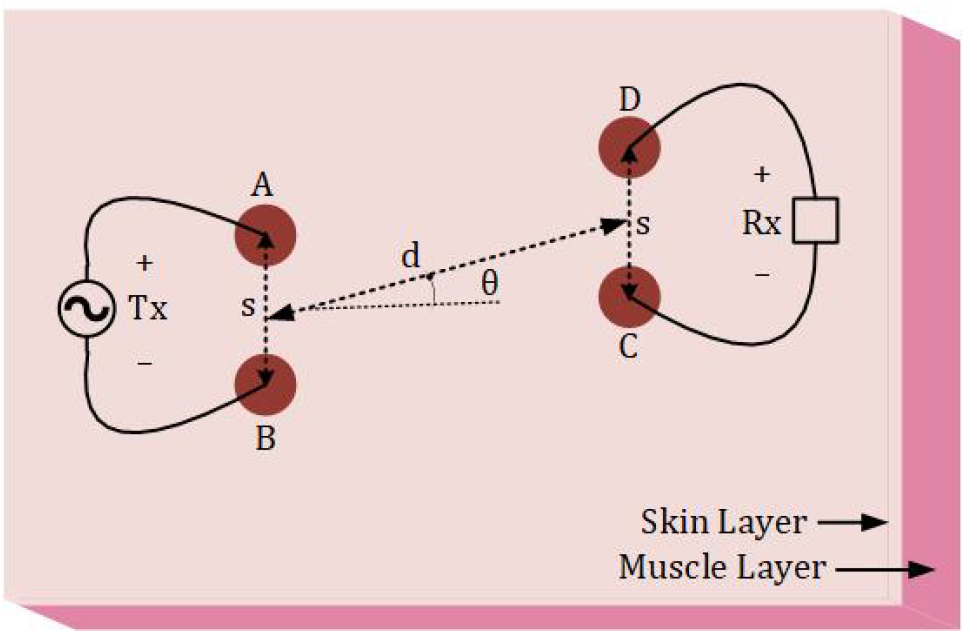
A pictorial representation of balanced Galvanic HBC with transmitting and receiving electrode separation s, channel length d and receiver’s angular position *θ* with respect to Tx.

In Voltage-Mode Human Body Communication [8] [17], the impedance provided by the return path capacitances are much higher than the impedance of the body in the signal path. So, the loading effect at the receiver side, due to the presence of return path capacitances is negligible. Therefore, considering negligible loading effect, the transfer function of the balanced galvanic HBC channel can be expressed with the help of Eq. 57 of **Appendix B**, by replacing impedance *Z*_0_ with *Z*_0*muscle*_ and impedance *Z*_*T*_ with (*Z*_*skin*_ + *Z*_*T muscle*_), such that

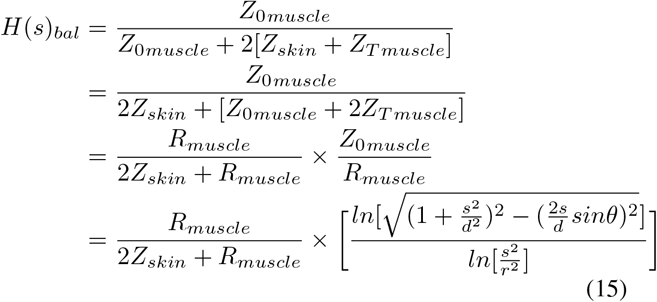

where, 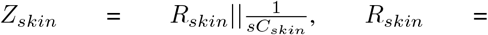 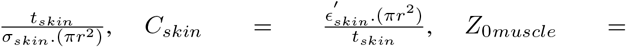 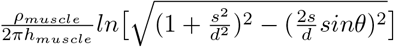 and 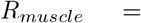 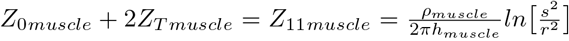. The first factor of Eq. 15 is coming from the voltage division between the skin and muscle layer, whereas the second factor is coming from the geometry dependant loss due to the relative position of the receiving electrodes with respect to the transmitting electrodes. The transfer function Eq. 15 can be simplified further into Eq. 16, by replacing *Z*_*skin*_ with 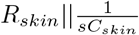, such that

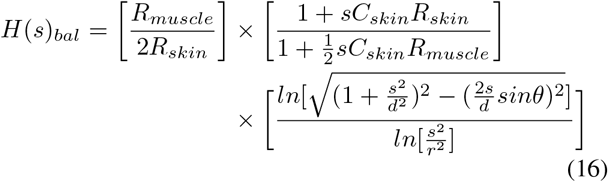

where, we consider *R*_*skin*_ ≫ *R*_*muscle*_ and 2*R*_*skin*_ + *R*_*muscle*_ ≃ 2*R*_*skin*_. Now using Eq. 16 a balanced galvanic HBC channel loss can be represented as

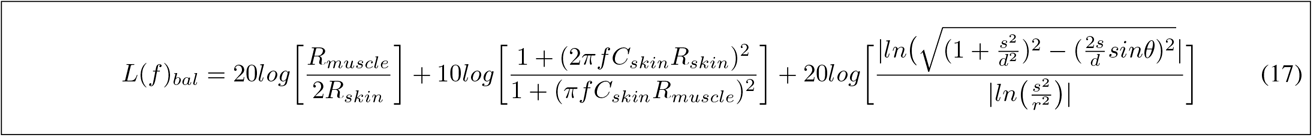

The loss expression *L*(*f*)_*bal*_, represented by Eq. 17 contains three different dependencies. The first one is due to the resistive division of the signal between skin and muscle layer at low frequency range which is mostly insensitive to the frequency. The second one is the frequency dependent loss which becomes effective at afrequency higher than a few kHz. The last one represents the dependency of the channel loss on channel length, electrode separation and the angular position of the receiver which can be called the loss due to the channel geometry and shape. The last part of the loss expression also tells the increases in channel loss as *d* and 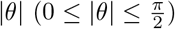 increases which is due to the reduction of impedance *Z*_0*muscle*_ in the proposed model.

To validate the loss expression, the channel loss obtained using Eq. 17 has been compared to the channel loss obtained from HFSS simulation. The frequency dependent parameters such as conductivity and permittivity for skin and muscle tissue layers are obtained from [10] and [11]. The comparative plots in Fig. 8(a) and Fig. 8(b) show that the proposed model is able to find the galvanic loss on a planner structure with an error margin less than 5 percent. The sharp voltage drop near the electrode edge in Fig. 8(b) which introduces mismatch between the simulation results and proposed model is due the the finite electrode size which can’t be neglected when the channel length is smaller than the electrode radius.

**Fig. 8:**
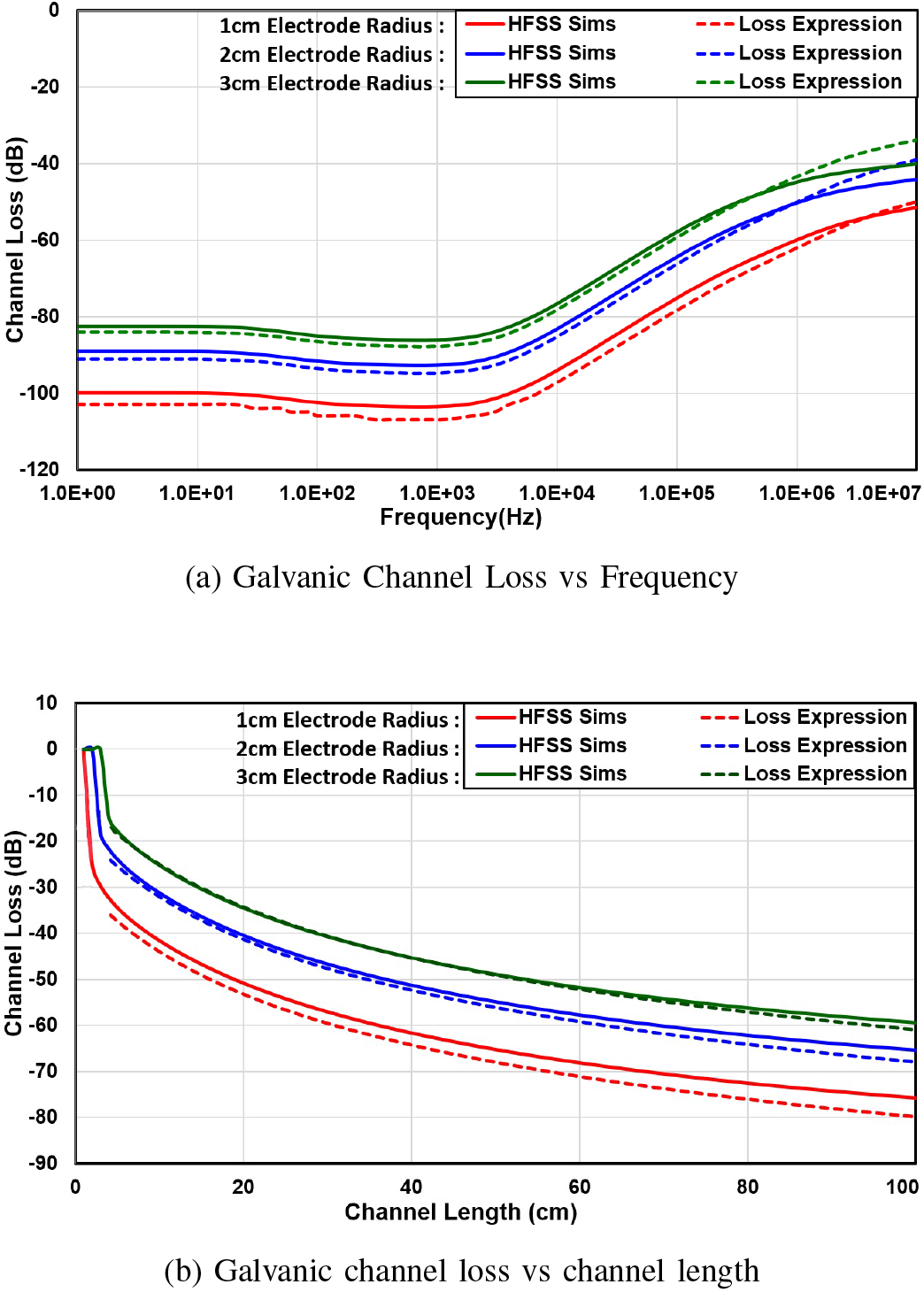
Comparison result of the simulation data with derived channel loss expression obtained from the proposed bio-physical model. (a) Represents the comparison plot of channel loss vs frequency for three different electrode sizes for a fixed 50cm channel length, (b) Represents the comparison plot of channel loss vs channel length for three different electrode sizes at a fixed 1 00kHz f requency o f operation;

## VI. Saturation of galvanic channel loss for an asymmetric or unbalanced channel

In the previous section, we have found the galvanic channel loss considering a balanced or symmetric HBC channel. But in most cases, the balanced nature of the HBC channel is not preserved due to the mismatch in the return path capacitances at the transmitter and receiver side. Eq. 15 shows, with balanced scenario the received voltage is proportional to the muscle impedance *Z*_0*muscle*_. At longer channel length *d* and higher *θ*, *Z*_0*muscle*_ tends to zero leading very small or almost zero received voltage. But in case of the unbalanced channel, due to the signal flow from the transmitter to the receiver through the return path capacitances via Earth-ground, the received voltage saturates to a fixed value when *Z*_0*muscle*_ reduces to a certain number. In this section, we have found the limiting value of the channel loss for a very long asymmetric channel.

In case of a very long body channel with a complicated geometry, we can consider the electrode distances *r*_*AC*_, *r*_*AD*_, *r*_*BC*_ and *r*_*BD*_ shown in Fig. 7 are approximately equal to the effective channel length *d* (≫*s*) which changes Eq. 12, Eq. 13 and Eq. 14 into Eq. 18, Eq. 19 and Eq. 20 respectively.

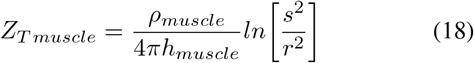

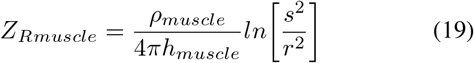

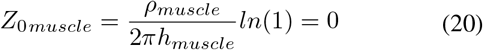

Now, with extremely low or zero *Z*_0*muscle*_ impedance, the received potential becomes a function of input common-mode potential 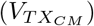 and the mismatch in the return path capacitances present at the transmitting and receiving side as derived in **Appendix B**. Therefore, the transfer function of a Galvanic HBC with an unbalanced channel can be represented using Eq. 61 of **Appendix B** by replacing *Z*_*R*_ with (*Z*_*skin*_ + *Z*_*Rmuscle*_).

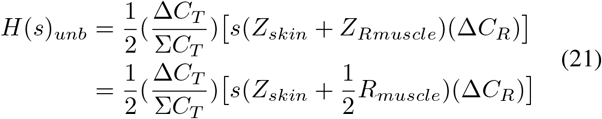

where, we have considered Δ*C*_*T*_ (= *C*_*ret*(*Tx*−)_−*C*_*ret*(*Tx*+)_) is the mismatch in the return path capacitance and Σ*C*_*T*_ (= *C*_*ret*(*Tx*+)_ + *C*_*ret*(*Tx*−)_) is the total return path capacitance at transmitter side; Δ*C*_*R*_(= *C*_*ret*(*Rx*−)_ − *C*_*ret*(*Rx*+)_) is the mismatch in the return path capacitance at receiver side; 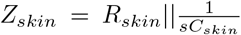, 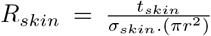, 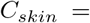 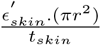 and 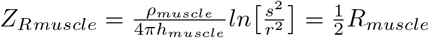.

The transfer function represented by Eq. 21 can be simplified further into Eq. 22, by replacing *Z*_*skin*_ with 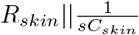, such as

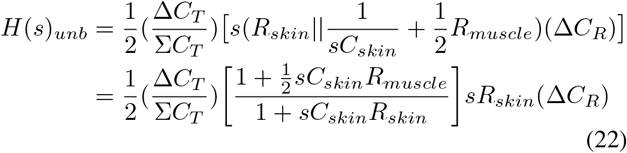

where, we consider 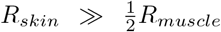 and 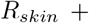 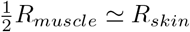.

In the sub-MHz frequency range it is seen that 1 ≪ *sC*_*skin*_*R*_*skin*_ and 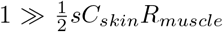 [10] [11]. Therefore in this frequency range the transfer function can be further reduces to Eq. 23.

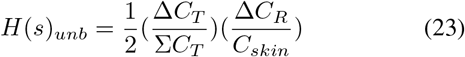

Using Eq. 23, channel loss can be represented as

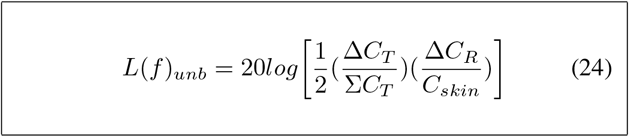

From Eq. 24 it is seen that, **Galvanic loss for a long and unbalanced channel is a function of mismatch in return path capacitances**. There has to be a mismatch in return capacitance at both the transmitting and receiving side to obtain a non zero received voltage. Mismatch at the transmitter side introduces a common-mode (CM) signal from the differential mode input. That common-mode(CM) gets converted again to a differential mode at the receiver side due to the mismatch present in the receiver electrodes. More the mismatch product lesser the signal loss is.

## VII. Extending the Concept for Cylindrical Structure

In the previous two sections we have formulated the galvanic HBC channel loss considering a planner structure of skin and muscle layer. But due to the complex structure of a human body, a planner structure is not enough to understand the signal loss around the body. Here in this section, we have extended the concept of the planner structure to a cylindrical body structure. We start with the E-field around the body for different type of excitations and validate the concept of planner structure what we derived in the previous sections.

Fig. 9 shows the strength of the Electric Field around a cylindrical structure made of skin and muscle tissue layer for three different excitation types. Fig. 9(a) shows the E-field pattern around the cylindrical structure with a transmitter connected capacitively. From the field pattern, it is seen that the E-filed is available everywhere around the body which ensures relatively less variant channel loss for capacitive HBC. Fig. 9(b) shows the E-field pattern of a balanced galvanic HBC. Here the E-field is restricted to a smaller region and dies out rapidly as distance from the transmitter increases. That is why channel loss in balanced galvanic HBC increases with channel length. The surface potential around the transmitting region shows equal and out of phase distribution of input voltage to the transmitting electrode pair. Equal and out of phased input voltage introduce zero common-mode voltage at the transmitting side and that confined the field within a small region. Fig. 9(c) shows the field pattern of an unbalanced galvanic HBC. Here, surface potential around the transmitting side shows unequal voltage distribution across the transmitting electrode pair. The unequal voltages at the transmitting electrodes introduce common-mode potential and that common-mode potential raises the overall body potential which can be picked up anywhere around the body. Because of the common-mode potential, the E-Filed is not restricted to a smaller region like the balanced transmission shown in Fig. 9(b). From the E-field pattern, it can also be concluded that **at longer channel length an unbalanced Galvanic HBC acts like capacitive HBC** where the loss is independent of channel length. At shorter channel length, the received voltage is due to the differential mode voltage as well as common-mode voltage but the voltage due to the differential mode dominates, and that makes the received voltage a function of channel length. At longer channel length differential voltage becomes very low and only the common-mode voltage contributes to the received potential.

**Fig. 9:**
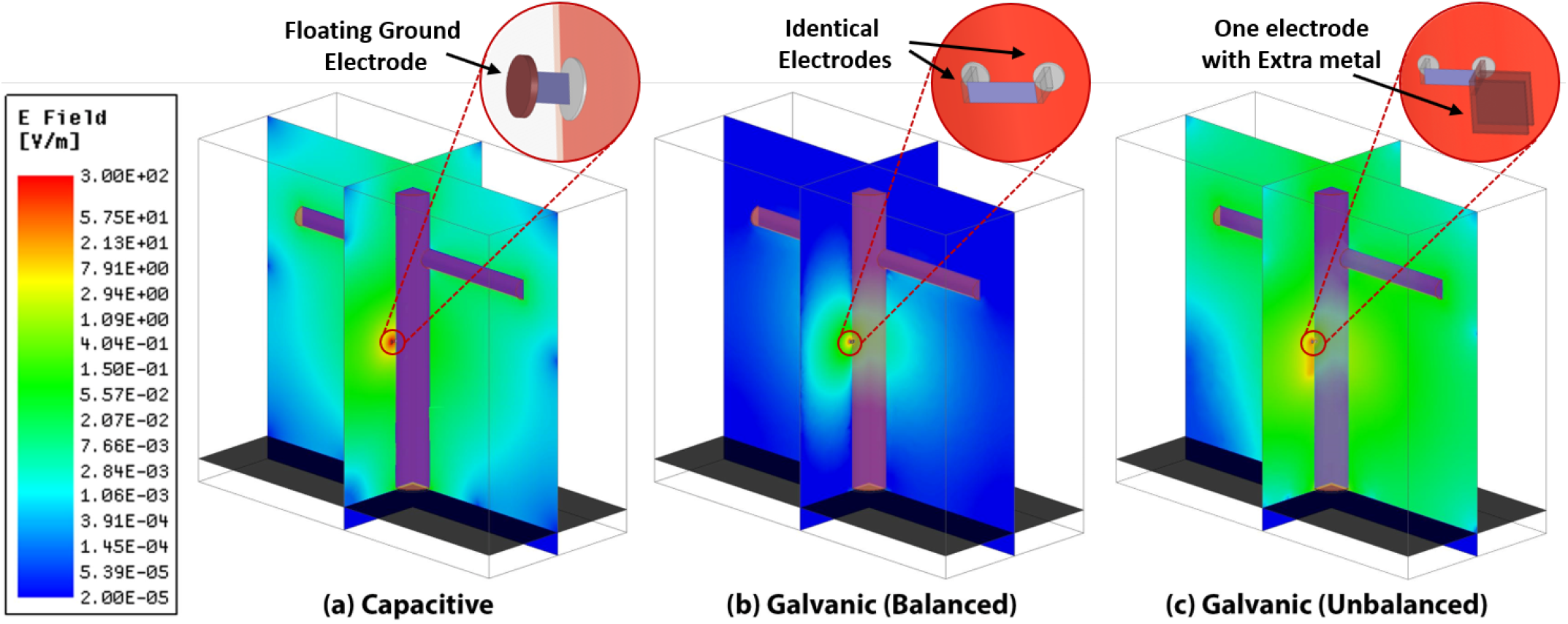
E-Field around the Human body with (a) Capacitive Excitation, (b) Balanced Galvanic Excitation and (c) Unbalanced Galvanic Excitation.

Fig. 10(a) shows the galvanic channel loss with respect to the channel length. In case of a balanced channel, the loss increases as channel length increases while in case of an unbalanced channel the loss increases with channel length and saturates to a fixed value for a longer channel length. The saturation value is determined by the mismatch present at the receiver and transmitter side as expressed by the Eq. 24.

**Fig. 10:**
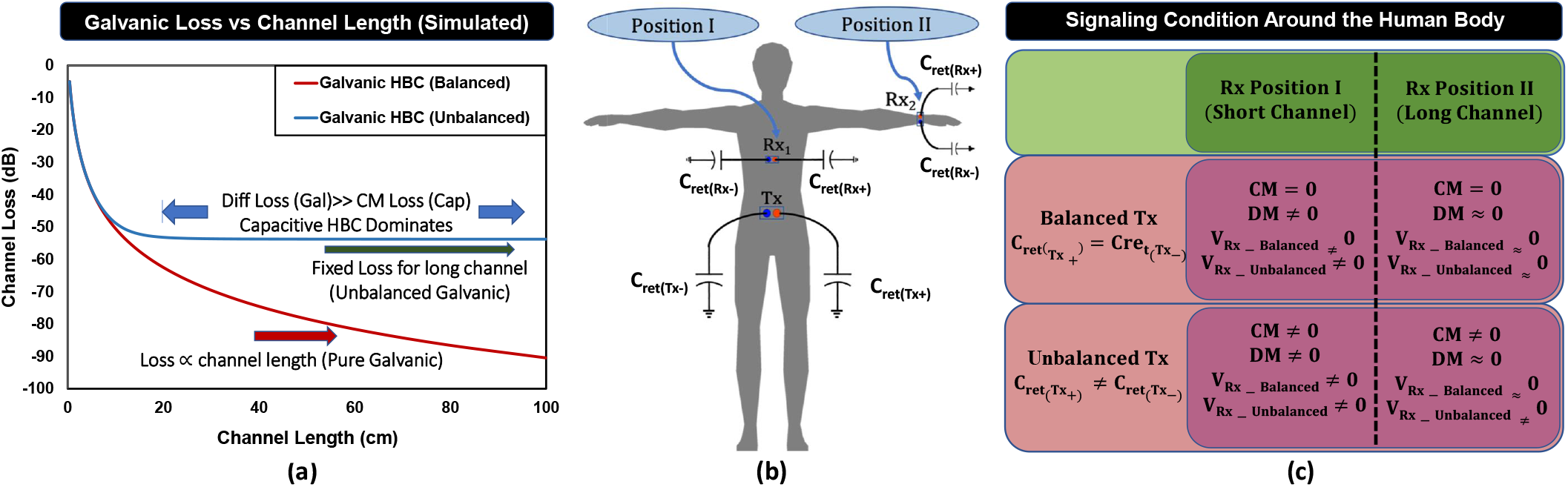
(a) The Simulation results of the distance dependent loss with Balanced or Unbalanced Tx-Rx pair; (b) and (c) shows the signaling condition around the human body in Galvanic arrangement considering short and long channel length.

Fig. 10(b) shows different types of signaling conditions depending on the nature of the channel. The channel between the transmitter and the first receiver *Rx*_1_ is considered to be a short channel and the distance between the transmitter and second receiver *Rx*_2_ is considered to be a long channel. The nature of the received signal depending on the balanced or unbalanced conditions are shown in Fig. 10(c).

## VIII. Measurement

In this section, we perform galvanic channel measurements to ascertain the validity of the proposed model.

### A. Measurement Setup

The channel loss measurement setup with Transmitter and Receiver connected in Galvanic arrangement is shown in Fig. 11. A ‘Velleman Instruments’ make battery-operated portable signal generator which generates frequency up to 1 MHz is used to generate a sinusoidal signal at 400KHz. At the receiver side, another battery-operated portable spectrum analyzer from RF Explorer is used to capture the signal. To reduce the wireless coupling between Tx and Rx, the electrodes are kept as close as to the signal generating or receiving port of those instruments. The connection between the signal generator and receiver with electrodes are made using coaxial SMA connectors to keep the signal shielded from outside noise. Circularly shaped copper metal with radius 1*cm* is used as an electrode. Each electrode pair of Tx and Rx are kept 5*cm* (center to center) apart from each other during the entire measurement process.

**Fig. 11:**
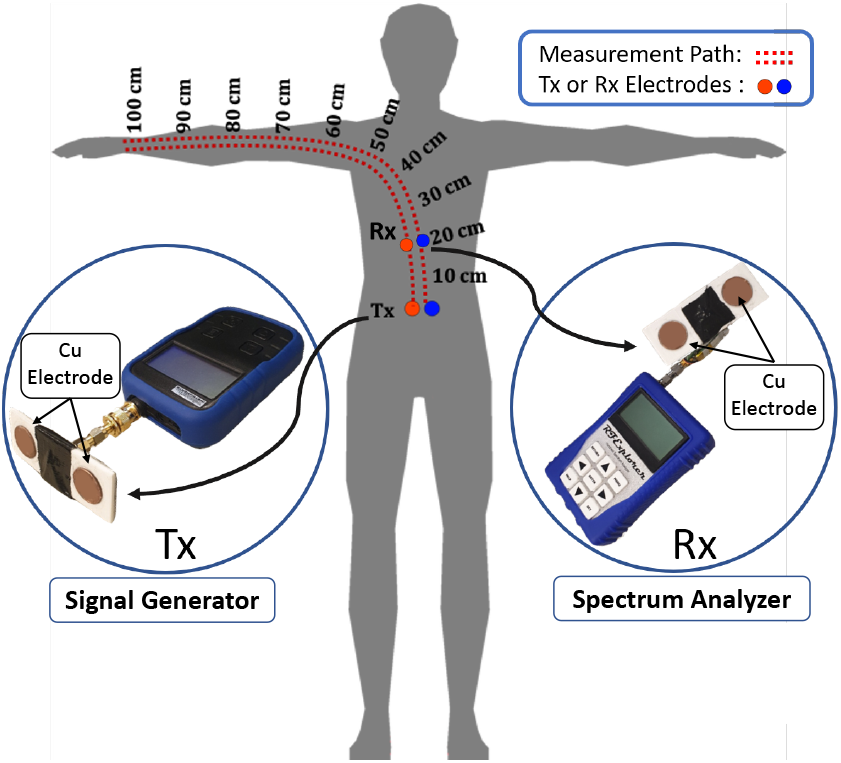
Diagram showing the position of Tx (fixed at waist) and Rx (moved along the dotted path) during the measurement.

### B. High Impedance Termination

The portable spectrum analyzer what has been used here as Rx has 50Ω input impedance which makes a low impedance termination at the receiver side. As described in [8], the low-frequency flat band response is only possible if the channel has a high impedance capacitive termination. A voltage buffer after the receiver electrodes provides the high impedance termination of the HBC channel. A 3.7V LiPo battery is used to provide supply to the Op-amp. A resistive divider is used to generate the input bias required to couple the HBC signal Fig. 12.

**Fig. 12:**
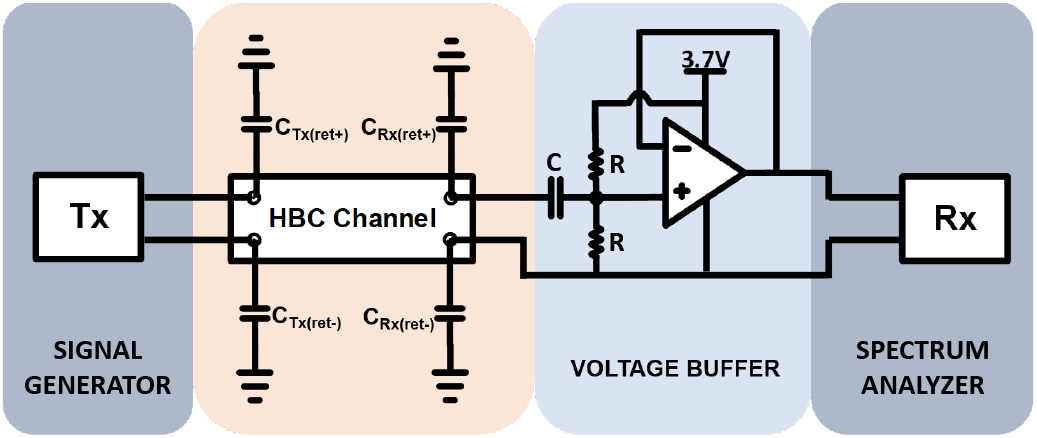
The measurement circuit diagram.

### C. Measurement Results

The path loss measurement is performed on multiple subjects with channel length starting from 2cm to 100cm. The Galvanic transmitter is kept on the waist part of the body with both the electrodes touching the skin. The signal is probed using the receiver along a path as shown in Fig. 11. The measurement results on three different subjects with three runs each are shown in Fig. 13. It is seen that, for a shorter channel length, loss increases with length and saturates to a constant value for channel length longer than 50cm. The possible explanation of this kind of measurement results can be traced back to the selection of the transmitter and receiver module. The portable instruments that we used during the measurement, mostly have a metallic casing which is connected to the ground of the device. The coupling between the metallic casing and Earth-Ground introduces bigger return capacitance at the negative plate of the Tx and Rx. While using those instruments in Galvanic arrangement, the positive electrode encounters smaller electrode to Earth-ground capacitance compare to the negative electrode i.e. *C*_*ret*(*Tx*+)_ ≪ *C*_*ret*(*Tx*−)_ and *C*_*ret*(*Rx*+)_ ≪ *C*_*ret*(*Rx*−)_. In case of our measurement setup, an independent experiment shows that, the extra capacitance to the negative plate of Galvanic devices are around 5pF-10pF which is 10 to 20 times more compare to the capacitance present in the positive electrode. The higher return path capacitance in one of the electrodes leads to an asymmetric scenario which helps to hold a fixed loss at longer channel length. Due to the unavailability of a portable as well as symmetric transceiver, it is difficult to measure true symmetric galvanic loss. However, the measurement result proves that, due to the presence of capacitive mismatch a Galvanic HBC channel starts to behave like a capacitive HBC channel at longer channel length. This Galvanic HBC to Capacitive HBC transform theory due to the mismatch has been discussed first time in this paper.

**Fig. 13:**
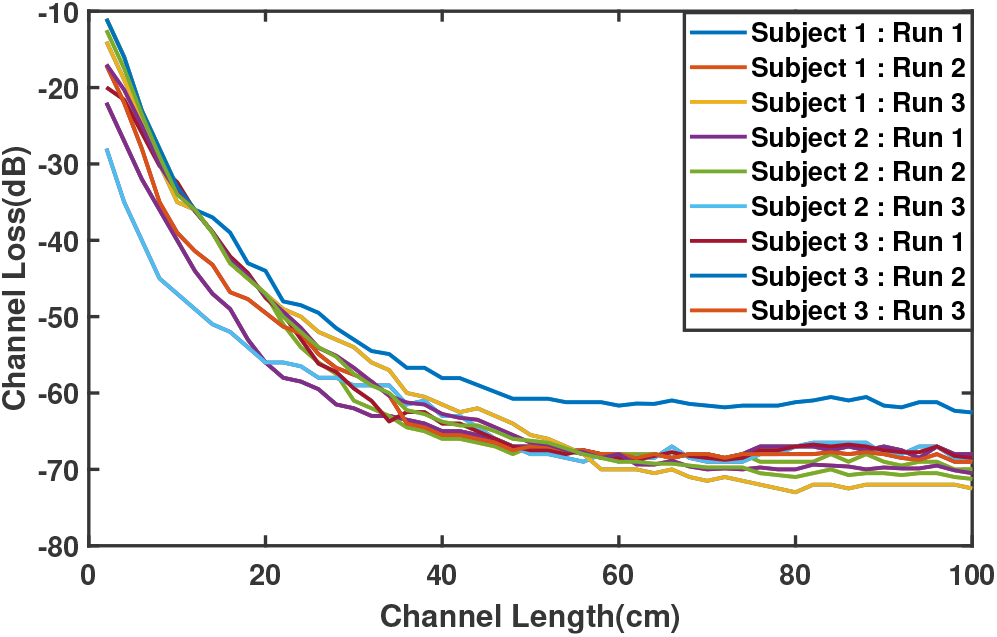
Measurement results of Galvanic HBC channel loss on three different subjects with three runs on each.

## IX. Implementation of Capacitive HBC model from the proposed Galvanic HBC model

In capacitive HBC, only one of the electrodes of transmitter/receiver is placed on the body whereas the other one is kept floating. Therefore, while transforming the Galvanic Bio-physical model to a capacitive bio-physical model, the skin capacitances and resistances which were connected to the negative plate of the Tx/Rx have to be neglected. Also the positive electrode of the transmitter and receiver get the extra body capacitance added to it compare to the negative plate. The transformation of an Galvanic Bio-physical model to a capacitive Bio-physical one is shown in Fir.14. First, we omitted the skin and muscle impedance for the negative plates leaving only skin-muscle-skin as a coupling medium between the positive Tx and Rx electrodes. Unlike galvanic HBC, in case of capacitive HBC the electrodes are highly unbalanced/asymmetric because of the body contact to the positive electrode only. The capacitance present to each of the electrode in capacitive connection can be expressed by Eq. 25–Eq. 28.

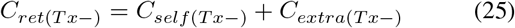

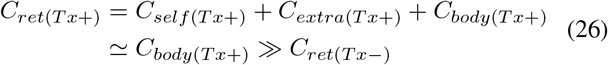

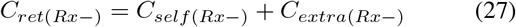

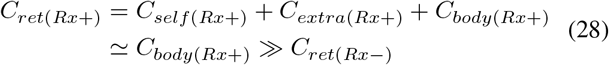

here, we approximated the total capacitance present at the positive electrode to the effective body capacitance as body capacitance is much higher than *C*_*self*_ and *C*_*extra*_ [8], [16]. Now neglecting the skin and muscle impedances (=*Z*_*skin*(*Tx*+)_ + *Z*_*muscle*_ + *Z*_*skin*(*Rx*+)_) which are much lower than the impedance provided by the return path capacitance, the model can be further reduced to a simplified one shown at the end of Fig. 14. In the simplified model the total body capacitance present in both the positive electrodes are combined to *C*_*body*_ such that

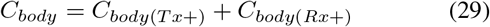

Finally, using the network as implemented at the end of Fig. 14, the transfer function of capacitive HBC can be expressed by Eq. 30

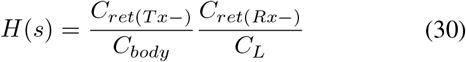

where, *C*_*L*_ is the effective load capacitance present between the receiver electrodes.

**Fig. 14:**
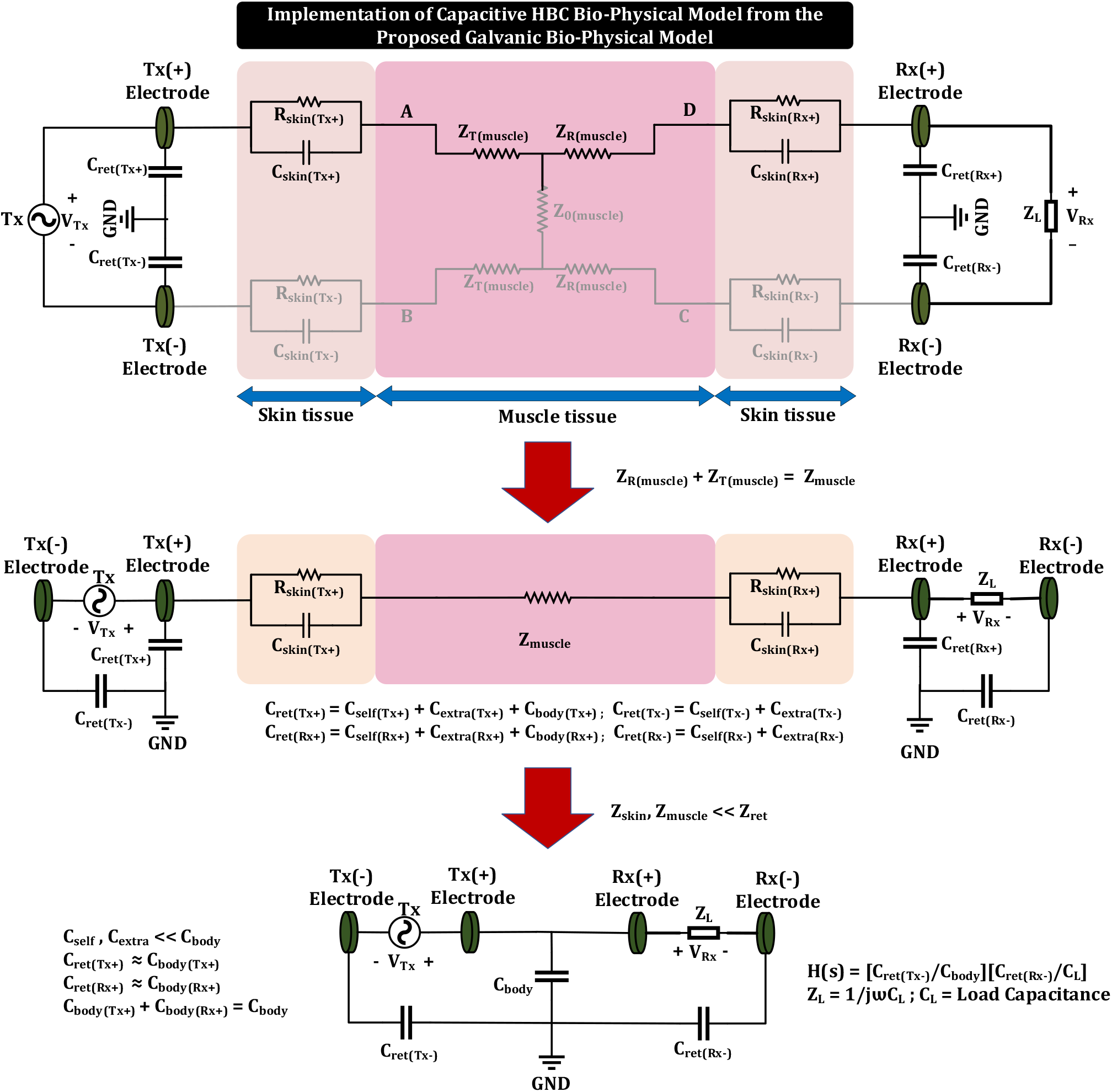
Implementation of Capacitive HBC Bio-physical model from the proposed Galvanic bio-physical model

## X. Conclusion

This paper characterizes the galvanic HBC channel in an Electro-Quasistatic frequency regime considering skin and muscle tissue layer as an in-body signal path along with four return path capacitances that form the outer body signal path. A unifying biophysical model has been proposed for the first time to capture the effects of in body and outer body signal path on channel loss. FEM simulation result of a planner structure of skin and muscle tissue layer is used to understand the behavior of the channel loss under balanced and unbalanced channel termination. Balanced or unbalanced termination of HBC channel arises due to the mismatch in return path capacitance at the transmitter and receiver side. It is also seen that the transmitted signal in galvanic HBC contains a non-zero common-mode(CM) due to the mismatch in the return path capacitance at the transmitter side. Due to the presence of return path capacitance to each of the electrodes, the signal appearing on the transmitting electrodes are completely out of phase with respect to the Earth-ground and the absolute amplitude of the signals on the electrodes are determined by the capacitance present at that electrode node. Unequal voltages to the transmitting electrodes in presence of any input mismatch or asymmetry, introduces a differential mode(DM) signaling with non-zero common-mode(CM). The differential signal reduces as the channel length increases, whereas the common-mode signal is independent of the channel length and propagate around the body with a fixed amount of loss. At the receiver end, if it is balanced i.e. both the electrodes contain equal return path capacitance, the receiver is only capable of receiving the differential signal available to its electrodes. In case of an unbalanced receiver i.e. with unequal return path capacitance at the receiver electrodes, it is capable of receiving a signal from the differential mode as well as common mode. At receiver end common mode to differential more conversion happens due to the unequal loading at the receiving electrodes. As the differential mode signal reduces with distance, the position of the receiver with respect to the transmitter (i.e. channel length) and the mismatch factor of both the transmitter and receiver decide the overall received signal level. At very short channel length the received signal is mostly due to the differential signal and at a longer channel length, the received signal is mostly due to the common-mode to differential mode conversion in case of an unbalanced channel where the channel behavior is dominated by the capacitive HBC, in spite of Galvanic excitation and termination.

## Notes

### Competing Interest Statement

The authors have declared no competing interest.

## References

[1] T. G. Zimmerman, “Personal Area Networks: Near-field intrabody communication,” IBM Systems Journal, vol. 35, no. 3.4, pp. 609–617, 1996.

[2] X. Wang, J. Shi, L. Xu, and J. Wang, “A Wideband Miniaturized Implantable Antenna for Biomedical Application at HBC Band,” in 2018 Cross Strait Quad-Regional Radio Science and Wireless Technology Conference (CSQRWC), Jul. 2018, pp. 1–3.

[3] N. Fahier and W. Fang, “An HBC-based continuous bio-potential system monitoring using 30mhz OOK modulation,” in 2017 IEEE Biomedical Circuits and Systems Conference (BioCAS), Oct. 2017, pp. 1–4.

[4] F. Koshiji, R. Urushidate, and K. Koshiji, “Biomedical signal transmission using human body communication,” in 2017 IEEE 36th International Performance Computing and Communications Conference (IPCCC), Dec. 2017, pp. 1–2.

[5] J. Wang, T. Fujiwara, T. Kato, and D. Anzai, “Wearable ECG Based on Impulse-Radio-Type Human Body Communication,” IEEE Transactions on Biomedical Engineering, vol. 63, no. 9, pp. 1887–1894, Sep. 2016.

[6] D. Das, S. Maity, B. Chatterjee, and S. Sen, “Enabling Covert Body Area Network using Electro-Quasistatic Human Body Communication,” Scientific Reports, vol. 9, no. 1, pp. 1–14, Mar. 2019. [Online]. Available: https://www.nature.com/articles/s41598-018-38303-x

[7] M. Hessar, V. Iyer, and S. Gollakota, “Enabling On-body Transmissions with Commodity Devices,” in Proceedings of the 2016 ACM International Joint Conference on Pervasive and Ubiquitous Computing, ser. UbiComp ‘16. New York, NY, USA: ACM, 2016, pp. 1100–1111, event-place: Heidelberg, Germany. [Online]. Available: http://doi.acm.org/10.1145/2971648.2971682

[8] S. Maity, M. He, M. Nath, D. Das, B. Chatterjee, and S. Sen, “Bio-Physical Modeling, Characterization, and Optimization of Electro-Quasistatic Human Body Communication,” IEEE Transactions on Biomedical Engineering, vol. 66, no. 6, pp. 1791–1802, Jun. 2019.

[9] M. S. Wegmueller, M. Oberle, N. Felber, N. Kuster, and W. Fichtner, “Signal Transmission by Galvanic Coupling Through the Human Body,” IEEE Transactions on Instrumentation and Measurement, vol. 59, no. 4, pp. 963–969, Apr. 2010.

[10] “Tissue Frequency Chart IT’IS Foundation.” [Online]. Available: https://itis.swiss/virtual-population/tissueproperties/database/tissue-frequency-chart/

[11] S. Gabriel, R. W. Lau, and C. Gabriel, “The dielectric properties of biological tissues: II. Measurements in the frequency range 10 Hz to 20 GHz,” Physics in Medicine and Biology, vol. 41, no. 11, pp. 2251–2269, Nov. 1996.

[12] M. S. Wegmueller, A. Kuhn, J. Froehlich, M. Oberle, N. Felber, N. Kuster, and W. Fichtner, “An attempt to model the human body as a communication channel,” IEEE Transactions on Biomedical Engineering, vol. 54, no. 10, pp. 1851–1857, 2007.

[13] K. Ito and Y. Hotta, “Signal path loss simulation of human arm for galvanic coupling intra-body communication using circuit and finite element method models,” in 2015 IEEE Twelfth International Symposium on Autonomous Decentralized Systems, 2015, pp. 230–235.

[14] X. M. Chen, P. U. Mak, S. H. Pun, C. Lam, U. K. Che, J. W. Li, Y. M. Gao, M. I. Vai, and M. Du, “Initial investigation of channel capacity for galvanic coupling human body communication,” in 2016 9th Biomedical Engineering International Conference (BMEiCON), Dec 2016, pp. 1–4.

[15] D. Ahmed, G. Fischer, and J. Kirchner, “Simulation-based models of the galvanic coupling intra-body communication,” in 2019 IEEE Sensors Applications Symposium (SAS), March 2019, pp. 1–6.

[16] M. Nath, S. Maity, and S. Sen, “Towards understanding the return path capacitance in capacitive human body communication,” IEEE Transactions on Circuits and Systems II: Express Briefs, pp. 1–1, 2019.

[17] S. Maity, D. Das, and S. Sen, “Wearable health monitoring using capacitive voltage-mode Human Body Communication,” in 2017 39th Annual International Conference of the IEEE Engineering in Medicine and Biology Society (EMBC), Jul. 2017, pp. 1–4.

