## Supplemental for "Bio-Physical Modeling of *Galvanic* Human Body Communication in Electro-Quasistatic Regime"

### Appendix A
### Two Port Equivalence of an Infinitely Stretched Conductor of Finite Thickness and Resistivity

An infinitely stretched conductive plane of resistivity $\rho$ and thickness $h$ is shown in the Fig.15 with a circular region of radius $r_A$ in the middle of the conductive surface. Current I is injected into the structure through the circular region of zero resistivity. Zero resistivity of the region ensures zero voltage drop inside the region. Now, to find the voltage profile on the surface of the conductor two concentric circles of radius *r* and *(r+dr)* are drawn with centers coinciding to the center of the region specified by radius $r_A$. If we assume the current is spreading out radially, the voltage difference between the annular ring specified by the circle of radius *r* and *(r+dr)* is

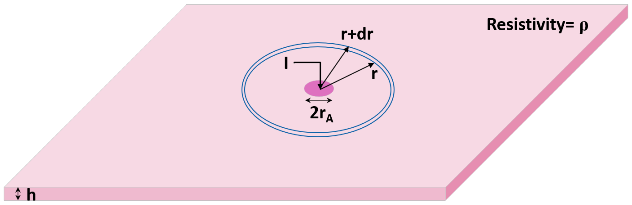


Fig. 15: An infinitely stretched conductive plane of thickness h and resistivity $\rho$. Current I is injected to the plane through a zero resistivity circular region of radius $r_A$.

$$dV = -\frac{\rho I}{2\pi r h} dr \tag{31}$$

Integrating Eq.31 over a radial distance $r_A$ to $r$ we have

$$V_r - V_{r_A} = \int_{r_A}^{r} -\frac{\rho I}{2\pi r h} dr = -\frac{\rho I}{2\pi h} ln\left[\frac{r}{r_A}\right] \tag{32}$$

where $V_{r_A}$ is the absolute potential of the circular region of radius $r_A$. Therefore, the voltage at a radial distance $r$ is a function of the radial distance $r$ and the absolute potential of the zero resistivity region where the current is injected Eq.33.

$$V_r = V_{r_A} - \frac{\rho I}{2\pi h} ln\left[\frac{r}{r_A}\right] \tag{33}$$

Now, considering another similar scenario where a current I is flowing through the same conductor from *Region A* to *Region B* as shown in the Fig.16. To find the voltage distribution on the conductive surface, we can consider the individual effect of the current of each zero resistivity region and use voltage superposition theorem to find the overall potential distribution at any arbitrary point on the conductive surface. Referring Eq.33, the potential of point P due to the injected current I at *Region A* can be written as

$$V_{P_A} = V_{r_A} - \frac{\rho I}{2\pi h} ln\left[\frac{r_{AP}}{r_A}\right] \tag{34}$$

where $V_{r_A}$ is the absolute potential of region A and $r_{AP}$ is the radial distance between the centre of the *region A* and *point P*. Similarly, assuming an injected current of $(-I)$ at *region B* the voltage at *point P* can be expressed as

$$V_{P_B} = V_{r_B} + \frac{\rho I}{2\pi h} ln\left[\frac{r_{AB}}{r_B}\right] \tag{35}$$

where $V_{r_B}$ is the absolute potential of region B and $r_{AB}$ is the radial distance between the centre of the *region B* and *point P*. Now, using voltage superposition theorem the overall potential at *point P* can be expressed as

$$\begin{aligned} V_P &= V_{P_A} + V_{P_B} \\ &= V_{r_A} + V_{r_B} + \frac{\rho I}{2\pi h} ln\left[\frac{r_A}{r_B}\frac{r_{BP}}{r_{AP}}\right] \end{aligned} \tag{36}$$

If point P in the Fig.16 approaches towards infinity the voltage $V_P$ approaches towards zero . Using this limiting condition i.e. $r_{AP} \to \infty$, $r_{AB} \to \infty$ leads $V_P \to 0$ in Eq.36 we have

$$V_P = V_{r_A} + V_{r_B} + \frac{I\rho}{2\pi h} ln(\frac{r_A}{r_B}) = 0 \tag{37}$$

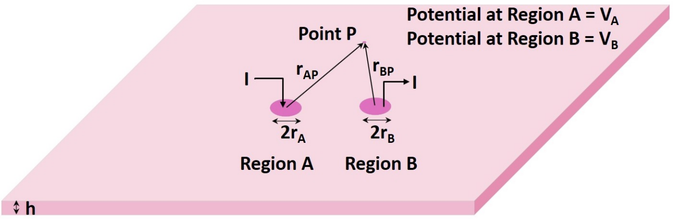


Fig. 16: An infinitely stretched conductive plane of thickness h and resistivity $\rho$ with I current flowing from region A to region B. Point P is an arbitrary point on the surface of the conductive plane at a radial distance $r_{AP}$ and $r_{BP}$ from the center of the region A and B respectively.

Therefore, combining Eq.36 and Eq.37, at any nearby point the potential distribution function of point P reduces to

$$V_P = \frac{I\rho}{2\pi h} ln(\frac{r_{BP}}{r_{AP}}) \tag{38}$$

The potential difference between Region A and Region B can now be found easily using Eq.38. To find the potential difference, let's consider a line $L$ connecting the center of the region A and center of the region B. The potential of the intersecting point of the line $L$ and perimeter of the Region A is the potential of region A, such that

$$V_A = \frac{I\rho}{2\pi h} ln(\frac{r_{AB} - r_A}{r_A}) \tag{39}$$

Similarly, the potential of the intersecting point of the line $L$ and perimeter of Region B is the potential of region B, such that

$$V_B = \frac{I\rho}{2\pi h} ln(\frac{r_B}{r_{AB} - r_B}) \tag{40}$$

Therefore, the voltage difference between Region A and Region B can be written as

$$\begin{aligned} V_{AB} &= V_A - V_B \\ &= \frac{I\rho}{2\pi h} ln(\frac{r_{AB} - r_A}{r_A}\frac{r_{AB} - r_B}{r_B}) \\ &\simeq \frac{I\rho}{2\pi h} ln(\frac{r_{AB}}{r_A}\frac{r_{AB}}{r_B}) \end{aligned} \tag{41}$$

where we have assumed that the value of the center to center radial distance between those electrodes is much higher than the value of the radius of the electrodes such that $r_{AB} \gg r_A$ and $r_{AB} \gg r_B$.

The potential distribution function i.e. Eq.38 can also be used to find the voltage difference between two arbitrary points on the surface of the conductor. Fig.17 shows the same conductor with four different circular regions. Current I flowing from Region A to Region B introducing a voltage difference between Region D and Region C. Again referring Eq.38, the voltage difference between Region D and Region C due to the current I can be defined as

$$\begin{aligned} V_{DC} &= V_D - V_C \\ &= \frac{I\rho}{2\pi h}\left[ln(\frac{r_{BD}-r_D}{r_{AD}-r_D}) - ln(\frac{r_{BC}-r_C}{r_{AC}-r_C})\right] \\ &\simeq \frac{I\rho}{2\pi h} ln\left[\frac{r_{AC}}{r_{AD}}\frac{r_{BD}}{r_{BC}}\right] \end{aligned} \tag{42}$$

where we have assumed the radial distances between those electrodes are much longer than the radius of the electrodes, such that $(r_{BD}, r_{AD}, r_{AC}, r_{BC}) \gg (r_A, r_B, r_C, r_D)$.

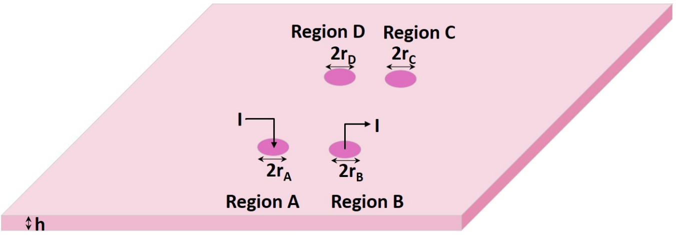


Fig. 17: An infinitely stretched conductive plane of thickness h and resistivity $\rho$ with I current flowing from region A to region B. Region D is an arbitrary point on the surface of the conductive plane at a radial distance $r_{AD}$ and $r_{BD}$ from the center of the region A and B respectively; whereas Region C is another arbitrary point on the surface of the conductive plane at a radial distance $r_{AC}$ and $r_{BC}$ from the center of the region A and B respectively.

Now, we can think the entire arrangement shown in Fig.17 as a two-port network, where Region A and Region B are used as an input port, whereas Region D and Region C act as an output port. Considering the concept of two-port network and using Eq.41 and Eq.42, representation of the conductor as a function of Z-impedance parameters is as follows

$$Z_{11} = \frac{V_{AB}}{I} = \frac{\rho}{2\pi h} ln\left[\frac{r_{AB}}{r_A}\frac{r_{AB}}{r_B}\right] \tag{43}$$

$$Z_{21} = \frac{V_{CD}}{I} = \frac{\rho}{2\pi h} ln\left[\frac{r_{AC}}{r_{AD}}\frac{r_{BD}}{r_{BC}}\right] \tag{44}$$

Applying the reciprocity theorem on the network the other two Z-parameters can be written as

$$Z_{22} = \frac{\rho}{2\pi h} ln\left[\frac{r_{CD}}{r_C}\frac{r_{CD}}{r_D}\right] \tag{45}$$

$$Z_{12} = Z_{21} = \frac{\rho}{2\pi h} ln\left[\frac{r_{AC}}{r_{AD}}\frac{r_{BD}}{r_{BC}}\right] \tag{46}$$

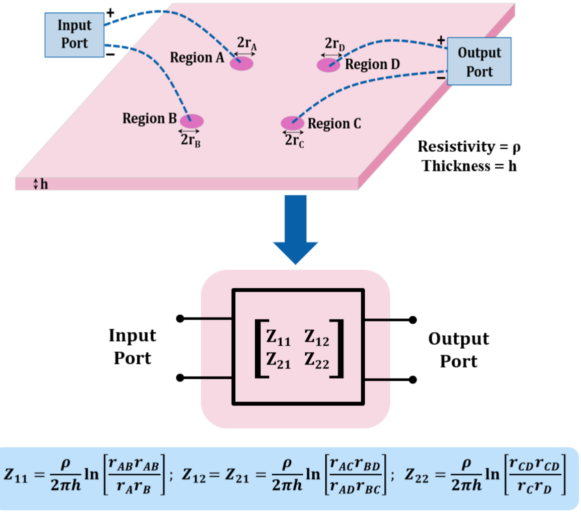


Fig. 18: Two port representation of an infinitely stretched conductive plane of thickness h and resistivity $\rho$ where Region A and Region B pair act as an input port and Region D and Region C pair act as an output port. The notation $r_i$ (i=A,B,C or D) denotes the radius of Region i whereas the notation $r_{ij}$ (i,j =A, B, C or D) denotes the center to center radial distance between Region i and Region j.

Therefore a planner conductive structure with a location for excitation and a location for the reception can be represented as a two-port network, where the network parameters $Z_{11}$, $Z_{12}$, $Z_{21}$ and $Z_{22}$ represent the electrical and geometric properties of the location of excitation and reception which define the relation between the input and output voltages.

### APPENDIX B
### TRANSFER FUNCTION OF THE PROPOSED GALVANIC HBC MODEL

The circuit shown in Fig.19 represents a proposed lumped circuit model of galvanic HBC, where the in body signal path has been formed by the impedance $Z_T$, $Z_0$ and $Z_R$. $Z_T$ and $Z_R$ represent the combined skin and muscle impedance at the transmitter and receiver side respectively; such that $Z_T = Z_{skin} + Z_{T_{(muscle)}}$ and $Z_R = Z_{skin} + Z_{R_{(muscle)}}$. The skin and muscle impedance are combined here to keep the analysis short and understandable. Capacitors $C_{T+}$, $C_{T-}$,$C_{R+}$ and $C_{R-}$ form outer body signal path due to the return path capacitor associated with each of the transmitting and receiving electrodes. The transmitter is connected between *Node A* and *Node B* with Node potential $V_{TX+}$ and $V_{TX-}$ respectively with respect to Earth-ground. The output is sensed across *Node D* and *Node C* which are at potential $V_{RX+}$ and $V_{RX-}$ respectively with respect to Earth-ground. Due to the mismatch present in the capacitors the circuit exhibits Differential Mode(DM) signal to Common Mode (CM) signal conversion or vice versa which is similar to the Differential

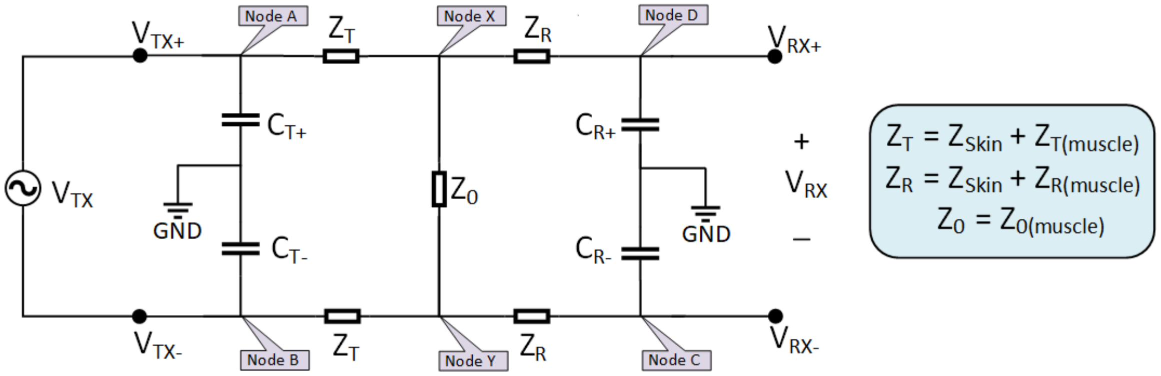


Fig. 19: Proposed Equivalent circuit of Galvanic HBC

Amplifier circuit. For better understanding, depending on the capacitance mismatch scenario, we have expressed the output voltage as a function of input Differential Mode(DM) voltage or input Common Mode (CM) voltage which itself is also a function of input Differential Mode(DM) voltage.

The capacitors connected to *Node A* and *Node B* with other terminal of the capacitors connected to Earth-ground set the absolute potential of *Node A* and *Node B* based on the capacitance of each of the capacitors. Applying voltage division between those capacitors the absolute potential of *Node A* and *Node B* can be expressed using Eq.47 and Eq.48.

$$V_{TX+} = \frac{C_{T-}}{C_{T+} + C_{T-}} V_{TX} = \left[\frac{C_{T-}}{\Sigma C_T}\right] V_{TX} \tag{47}$$

$$V_{TX-} = -\frac{C_{T+}}{C_{T+} + C_{T-}} V_{TX} = -\left[\frac{C_{T+}}{\Sigma C_T}\right] V_{TX} \tag{48}$$

where, $\Sigma C_T (= C_{T+} + C_{T-})$ is the total capacitance present at the input port. Using Eq.47 and Eq.48 the input common mode potential can be defined as

$$\begin{aligned} V_{TX_{CM}} &= \frac{1}{2}(V_{TX+} + V_{TX-}) \\ &= \frac{1}{2}\left[\frac{C_{T-} - C_{T+}}{\Sigma C_T}\right] V_{TX} \\ &= \frac{1}{2}\left(\frac{\Delta C_T}{\Sigma C_T}\right) V_{TX} \end{aligned} \tag{49}$$

where, $\Delta C_T (= C_{T-} - C_{T+})$ is the difference in the capacitance value present at the input port. Eq.49 shows how capacitance mismatch at the input port leads Differential Mode(DM) signal to non-zero Common Mode(CM) signal generation.

In HBC, the impedance of the return path capacitances are much higher than the impedance of the body. So, assuming the impedance due to the capacitors $C_{R+}$ and $C_{R-}$ much higher compare to the impedance $Z_0$ and using voltage superposition, the potential at *Node X* and *Node Y* can be expressed as

$$\begin{aligned} V_X &= \frac{Z_0 + Z_T}{Z_0 + 2Z_T} V_{TX+} + \frac{Z_T}{Z_0 + 2Z_T} V_{TX-} \\ &= \frac{Z_0 + Z_T}{Z_0 + 2Z_T}\left[\frac{C_{T-}}{\Sigma C_T}\right] V_{TX} - \frac{Z_T}{Z_0 + 2Z_T}\left[\frac{C_{T+}}{\Sigma C_T}\right] V_{TX} \\ &= \frac{1}{Z_0 + 2Z_T}\left[\frac{Z_0 C_{T-} + Z_T(C_{T-} - C_{T+})}{\Sigma C_T}\right] V_{TX} \\ &= \frac{1}{Z_0 + 2Z_T}\left[\frac{Z_0 C_{T-} + Z_T(\Delta C_T)}{\Sigma C_T}\right] V_{TX} \end{aligned} \tag{50}$$

$$\begin{aligned} V_Y &= \frac{Z_T}{Z_0 + 2Z_T} V_{TX+} + \frac{Z_0 + Z_T}{Z_0 + 2Z_T} V_{TX-} \\ &= \frac{Z_T}{Z_0 + 2Z_T}\left[\frac{C_{T-}}{\Sigma C_T}\right] V_{TX} - \frac{Z_0 + Z_T}{Z_0 + 2Z_T}\left[\frac{C_{T+}}{\Sigma C_T}\right] V_{TX} \\ &= -\frac{1}{Z_0 + 2Z_T}\left[\frac{Z_0 C_{T+} - Z_T(C_{T-} - C_{T+})}{\Sigma C_T}\right] V_{TX} \\ &= -\frac{1}{Z_0 + 2Z_T}\left[\frac{Z_0 C_{T+} - Z_T(\Delta C_T)}{\Sigma C_T}\right] V_{TX} \end{aligned} \tag{51}$$

Now, at the receiver side considering impedance division the potential at *Node D* and *Node C* can be expressed as

$$V_D = V_{RX+} = \frac{\frac{1}{sC_{R+}}}{Z_R + \frac{1}{sC_{R+}}} V_X = \left[\frac{1}{1 + sZ_R C_{R+}}\right] V_X \tag{52}$$

$$V_C = V_{RX-} = \frac{\frac{1}{sC_{R-}}}{Z_R + \frac{1}{sC_{R-}}} V_X = \left[\frac{1}{1 + sZ_R C_{R-}}\right] V_Y \tag{53}$$

Therefore the received voltage at the receiver side is

$$\begin{aligned} V_{RX} &= V_{RX+} - V_{RX-} \\ &= \left[\frac{1}{1 + sZ_R C_{R+}}\right] V_X - \left[\frac{1}{1 + sZ_R C_{R-}}\right] V_Y \end{aligned} \tag{54}$$

Using the voltage expression of $V_X$ and $V_Y$ from Eq.50 and Eq.51, the received voltage can be expressed as

$$\begin{aligned} V_{RX} = \frac{1}{Z_0 + 2Z_T}\Bigg[&\frac{Z_0 C_{T-} + Z_T(\Delta C_T)}{\Sigma C_T(1 + sZ_R C_{R+})} \\ &+ \frac{Z_0 C_{T+} - Z_T(\Delta C_T)}{\Sigma C_T(1 + sZ_R C_{R-})}\Bigg] V_{TX} \end{aligned} \tag{55}$$

Therefore, the transfer function of the network is

$$H(s) = \frac{V_{RX}}{V_{TX}} = \frac{1}{Z_0 + 2Z_T}\left[\frac{Z_0 C_{T-} + Z_T(\Delta C_T)}{\Sigma C_T(1 + sZ_R C_{R+})} + \frac{Z_0 C_{T+} - Z_T(\Delta C_T)}{\Sigma C_T(1 + sZ_R C_{R-})}\right] \quad (56)$$

Now, we are going to find the transfer function of the network considering two special cases. In the first case, we are going to find the transfer function for a fully symmetric or balanced network. Whereas in the other case, we will consider the effect of asymmetry or unbalance at the input and output port. In both cases, we will find the effect of $Z_0$ on the transfer function.

#### A. *Symmetric or Balanced Condition:*

A symmetric or balanced condition of the network occurs when $C_{T+} = C_{T-} = C_T$ and $C_{R+} = C_{R-} = C_R$ which leads $\Delta C_T = 0$, $\Delta C_R = 0$, $\Sigma C_T = 2C_T$ and $\Sigma C_R = 2C_R$. Under this symmetric condition the common mode potential at the input port becomes zero as $\Delta C_T = 0$ in Eq.49. The potential at the receiver side is also fully differential with zero common mode as $\Delta C_R = 0$. Using Eq.56 and considering symmetric condition the transfer function of the network becomes

$$\begin{aligned} H(s)_{Sym} &= \frac{1}{Z_0 + 2Z_T} \times \left[\frac{Z_0 C_T + Z_0 C_T}{2C_T(1 + sZ_R C_R)}\right] \\ &= \frac{Z_0}{Z_0 + 2Z_T} \times \left[\frac{1}{1 + sZ_R C_R}\right] \end{aligned} \quad (57)$$

With symmetric condition, Eq.57 shows, two factors mainly determine the transfer function of the network. The first factor represents the fraction of input voltage available across the $Z_0$ impedance assuming there is no loading effect due to the $C_{R+}$ and $C_{R-}$ capacitors. The second factor represents the fraction of the voltage across $Z_0$ available to the output port. Further consideration, $sZ_R C_R \ll 1$ reduces the transfer function of symmetric network to Eq.58

$$H(s)_{Sym} = \frac{Z_0}{Z_0 + 2Z_T} \quad (58)$$

As we can see from the Eq.57 or Eq.58 if $Z_0$ becomes negligibly small the received voltage reduces drastically and with $Z_0 = 0$ received voltage $V_{RX}$ becomes zero. Therefore in a symmetric condition with $Z_0 = 0$, there is no voltage at the output, but in presence of asymmetry, there is a chance of non zero received voltage which is discussed in the next section.

#### B. *Asymmetric or Unbalanced Condition:*

In presence of asymmetry i.e. $\Delta C_T \neq 0$, $\Delta C_R \neq 0$ the transfer function is already derived in Eq.56. But from Eq.56, it difficult to understand the behaviour of the transfer function as $Z_T$ and $Z_0$ change. Here we are considering two cases:

*1) Case I:* $Z_T(\Delta C) \ll Z_0 C_{T+}, Z_0 C_{T-}$
This scenario arises when the channel is relatively short. With this condition the transfer function becomes

$$H(s)_{Asym(short)} = \frac{Z_0}{Z_0 + 2Z_T}\left[\frac{C_{T-}}{\Sigma C_T(1 + sZ_R C_{R+})} + \frac{C_{T+}}{\Sigma C_T(1 + sZ_R C_{R-})}\right] \quad (59)$$

The transfer function is still a function of $Z_0$ and as $Z_0$ reduces the value of the transfer function reduces.

*2) Case II:* $Z_T(\Delta C) \gg Z_0 C_{T+}, Z_0 C_{T-}$
This scenario arises when the channel is relatively long. Therefore, for asymmetric long channel the transfer function reduces to

$$\begin{aligned} H(s)_{Asym(long)} &= \frac{Z_T}{Z_0 + 2Z_T}\left[\frac{(\Delta C_T)}{\Sigma C_T(1 + sZ_R C_{R+})} - \frac{(\Delta C_T)}{\Sigma C_T(1 + sZ_R C_{R-})}\right] \\ &= \frac{1}{2}\left(\frac{\Delta C_T}{\Sigma C_T}\right)\left[\frac{1}{1 + sZ_R C_{R+}} - \frac{1}{1 + sZ_R C_{R-}}\right] \end{aligned} \quad (60)$$

Here we also considered $Z_0 \ll Z_T$ which is obvious for a long channel. Now considering $sZ_R C_{R+} \ll 1$ and $sZ_R C_{R-} \ll 1$ which is due to the high impedance of return path capacitance compared to the body impedance, the transfer function for the asymmetric condition can be further reduced to

$$\begin{aligned} H(s)_{Asym} &= \frac{1}{2}\left(\frac{\Delta C_T}{\Sigma C_T}\right)\left[(1 - sZ_R C_{R+}) - (1 - sZ_R C_{R-})\right] \\ &= \frac{1}{2}\left(\frac{\Delta C_T}{\Sigma C_T}\right)\left[sZ_R(C_{R-} - C_{R+})\right] \\ &= \frac{1}{2}\left(\frac{\Delta C_T}{\Sigma C_T}\right)\left[sZ_R(\Delta C_R)\right] \end{aligned} \quad (61)$$

Therefore, the received potential for an asymmetric long channel is only a function of mismatch in the return path capacitance. Absence of $Z_0$ impedance in the transfer function makes the output voltage independent of channel length in asymmetric long channel scenario. The received potential can also be expressed as a function of common-mode input voltage which is defined by Eq.49 earlier.

$$\begin{aligned} V_{RX} &= \frac{1}{2}\left(\frac{\Delta C_T}{\Sigma C_T}\right)\left[sZ_R(\Delta C_R)\right]V_{TX} \\ &= \left[sZ_R(\Delta C_R)\right]V_{TX_{CM}} \end{aligned} \quad (62)$$

Eq.62 establishes the relation between the received signal($V_{RX}$) to the input common-mode ($V_{TX_{CM}}$) signal which is a function of input mismatch and input differential voltage ($V_{TX}$). For this asymmetric and long channel scenario, a non zero output is only possible if the mismatch is present at the input as well as at the output side. The input mismatch converts the Differential input voltage to the Common mode voltage due to the unequal voltage division across the capacitors at that port. At the output side, due to the mismatch in the capacitors the input Common mode signal is again converted to a differential mode signal due to the unequal voltage division at that port.

Thus, we can conclude that, for a symmetric or balanced scenario, the received potential is proportional to $Z_0$ impedance. In case of an asymmetric or unbalanced scenario, the received potential is a function of impedance $Z_0$ when the impedance $Z_0$ is comparatively large. At comparatively smaller $Z_0$ value the received potential becomes a function of capacitor mismatch of the input and output ports.
